# Site-specific phosphorylation of PSD-95 dynamically regulates the post-synaptic density as observed by phase separation

**DOI:** 10.1101/2021.03.03.433754

**Authors:** Maria Vistrup-Parry, Xudong Chen, Thea L. Johansen, Sofie Bach, Sara C. Buch-Larsen, Christian R. O. Bartling, Chenxue Ma, Louise S. Clemmensen, Michael L. Nielsen, Mingjie Zhang, Kristian Strømgaard

## Abstract

Postsynaptic density protein 95 is a key scaffolding protein in the postsynaptic density of excitatory glutamatergic neurons, organizing signaling complexes primarily via its three PSD-95/Discs-large/Zona occludens domains. PSD-95 is regulated by phosphorylation, but technical challenges have limited studies of the molecular details. Here, we genetically introduced site-specific phosphorylations in single, tandem and full-length PSD-95 and generated a total of 11 phosphorylated protein variants. We examined how these phosphorylations affected binding to known interaction partners and the impact on phase separation of PSD-95 complexes, and identified two new phosphorylation sites with opposing effects. Phosphorylation of Ser78 inhibited phase separation with the glutamate receptor subunit GluN2B and the auxiliary protein stargazin, whereas phosphorylation of Ser116 induced phase separation with stargazin only. Thus, by genetically introducing phosphoserine site-specifically and exploring the impact on phase separation, we have provided new insights into the regulation of PSD-95 by phosphorylation and the dynamics of the PSD.

**Graphical Abstract: Phosphorylation of PSD-95.**

A. Illustration of the interaction between the PDZ domains of PSD-95 and the NMDA receptor (NMDAR) (PDB: 4PE5) at the plasma membrane. PSD-95 interacts with the AMPA receptor (AMPAR) (PDB: 3KG2) through Stargazin (PDB: 5KBU). Using the software Ranch, a model of FL PSD-95 was generated using known structural domains of PSD-95 connected via a fully-flexible linker (PDZ1-2, PDB: 3GSL; PDZ3, PDB: 5JXB; SH3-GK, PDB: 1KJW).
B. Illustration of the effect of phosphorylation of PSD-95 on the formation of condensates at the PSD. PSD-95 can phase separate with receptors and intracellular proteins, and phosphorylation is can affect this interaction and lead to alterations, both negative and positive, in the formation of condensates.

## INTRODUCTION

The postsynaptic density protein 95 (PSD-95) is one of the most abundant proteins in the postsynaptic density (PSD) of excitatory glutamatergic neurons (Zhu et al., 2016). PSD-95 interacts with receptors, ion channels and intracellular proteins thereby organizing signaling complexes critical for synaptic development and transmission (Feng and Zhang, 2009, Kim and Sheng, 2004). At the plasma membrane, PSD-95 interacts with the glutamate receptor subtypes α-amino-hydroxy-5-methyl-4-isoxazolepropionic acid (AMPA) receptor through its auxiliary subunit stargazin (Stg) (Chen et al., 2000, El-Husseini et al., 2000, Nicoll et al., 2006) and directly with the C-terminal of the *N*-methyl-D-aspartate (NMDA) receptor (Niethammer et al., 1996, Kornau et al., 1995), amongst others, and with components of the PSD and signaling proteins like neuronal nitric oxide synthase (nNOS) (Sattler et al., 1999, Pedersen et al., 2014, Aarts et al., 2002). PSD-95 has been implicated in postsynaptic development, as it is one of the first proteins to cluster at synapses (Rao et al., 1998, Funke et al., 2005). Moreover, mutations in PSD-95 have been identified as associated with diseases such as stroke, as well as with intellectual disability, autism spectrum disorder and schizophrenia (Volk et al., 2015, Coley and Gao, 2018).

PSD-95 belongs to the family of membrane associated guanylate kinase (MAGUK) proteins and contains three PSD-95/Discs-large/Zona occludens (PDZ) domains along with an Src Homology 3 (SH3) and a guanylate kinase (GK) domain (Figure 1A) (Chen et al., 2015). PDZ domains are one of the largest classes of protein-protein interaction (PPI) modules, and the vast majority of PSD-95 mediated interactions are attributed to the three PDZ domains of PSD-95 (Ye and Zhang, 2013, Lee and Zheng, 2010). PDZ domains are comprised of approximately 90 amino acids arranged in a specific globular fold (Doyle et al., 1996), and more than 250 human proteins have been acknowledged to contain PDZ domains (Ye and Zhang, 2013). In the canonical binding mode PDZ domains bind the extreme carboxy-terminal of the protein interaction partner (Doyle et al., 1996, Kim et al., 1995, Chi et al., 2012), characterized by the consensus sequence S/T-X-Φ-OH, where X is any amino acid and Φ is a hydrophobic amino acid. Some PDZ domains although confer non-canonical binding of internal peptide motifs (Harris et al., 2001).

**Figure 1:**
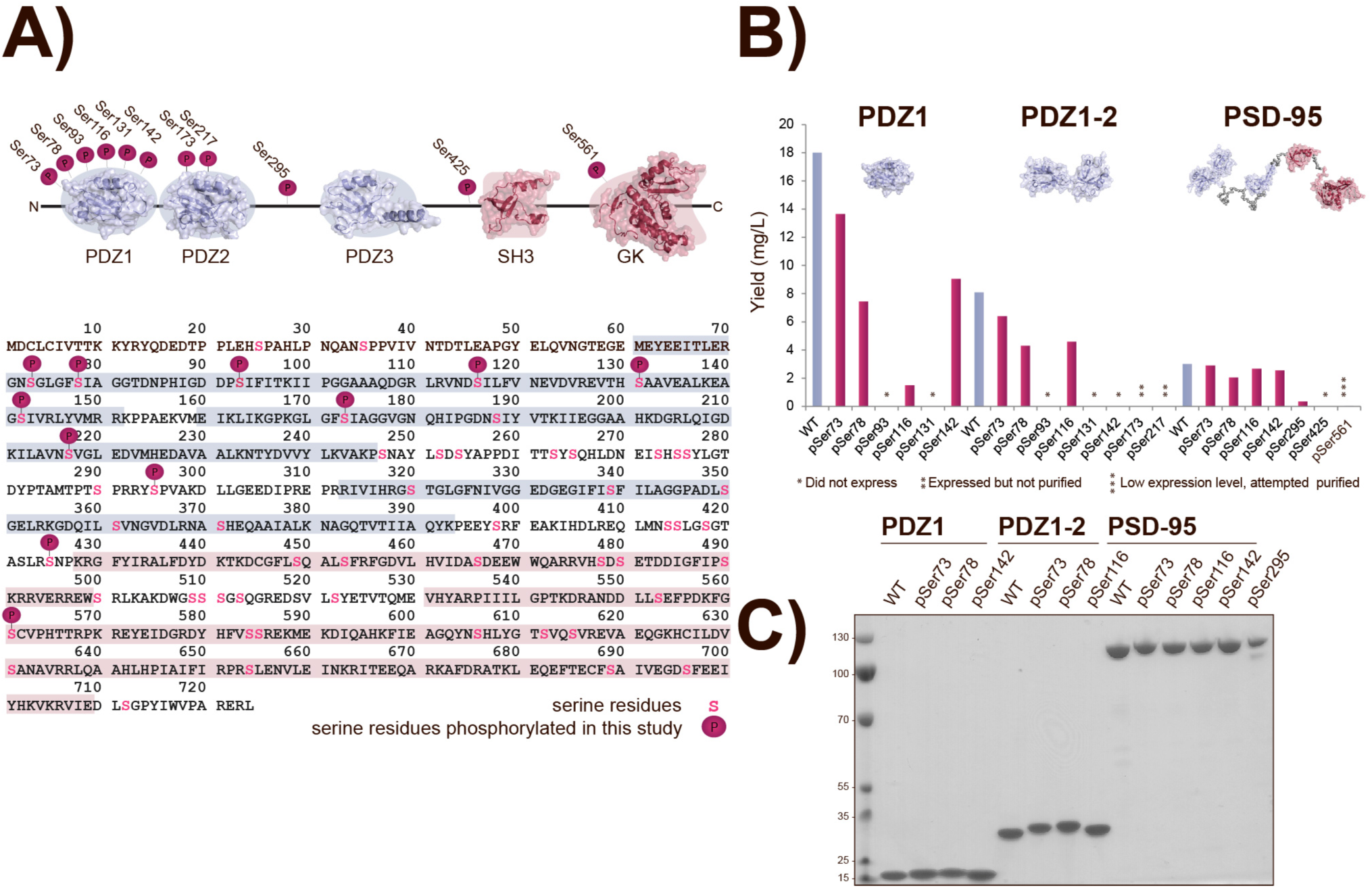
Site-specific phosphorylation of PSD-95. **A)** The organization of the five domains in PSD-95 (PDZ1-3, SH3 and GK) illustrated above the primary sequence of human FL PSD-95. All Ser residues are highlighted in pink, and the Ser residues attempted phosphorylated in this study highlighted by a phosphorylation icon. **B)** Histogram depicting all sites attempted phosphorylated and the yield (mg/L) of purification for purified proteins for all three protein forms. Proteins that did not express are indicated by a *, proteins that expressed but have not been purified by **, and proteins with low expression and attempted purified with ***. **C)** Coomassie stained gel of all purified pSer protein variants loaded alongside a protein ladder.

PPIs generally play essential roles in all aspects of cellular function by mediating signal transduction and metabolism, and due to many diseases being linked to malfunctioning signaling, PPIs have been explored as drug targets. In particular, the PDZ-mediated interactions between PSD-95, NMDA receptors and nNOS have been explored in the treatment of ischemic stroke, and two inhibitors of PSD-95, NA-1 (nerinetide) and AVLX-144, are currently in different stages of clinical trials (Hill et al., 2012, Bach et al., 2012).

The PSD was first visualized by electron microscopy (EM) as an electron-dense thickening below the plasma membrane (Gray, 1959). Since, the organization of the PSD has been shown to constitute multiple proteins in addition to PSD-95, which anchors proteins in the plasma membrane to the cytoskeleton (Sheng and Hoogenraad, 2007, Sheng and Kim, 2011, Chen et al., 2011). The dynamics of the PSD have been challenging to study, as the PSD is a protein-rich membrane-less compartment that assembles and disassembles continuously (Hyman et al., 2014, Zeng et al., 2016). Furthermore, synapses are dynamic and plastic, and hence, the morphology of two neighboring synapses can differ significantly (Nishiyama and Yasuda, 2015, Zeng et al., 2018). However, recently the PSD was reconstituted and subsequently studied using liquid-liquid phase separation (LLPS) (Zeng et al., 2018). Here, the reconstituted PSD, consisting of PSD-95, guanylate kinase-associated protein (GKAP), SH3 and multiple ankyrin repeat domain protein (Shank) and Homer protein homolog 3 (Homer3), was able to cluster subunits of the NMDA receptor, enrich synaptic enzymes and promote actin bundle formation. Hence this model system proved to be a highly promising approach for studying the interactions of the PSD (Zeng et al., 2018).

PSD-95 is vastly regulated by phosphorylation thereby affecting synaptic activity (Vallejo et al., 2016, Gardoni et al., 2006). The first PDZ domain (PDZ1) of PSD-95 contains six Ser residues, and as Ser is the most prevalent acceptor residue of protein phosphorylation, multiple of these residues have been identified by mass spectrometry (MS)-based approaches to be phosphorylated (Ballif et al., 2008, Xue et al., 2008). Amongst these Ser73 has been shown to be phosphorylated by CaMKIIα (Gardoni et al., 2006). Phosphorylation of PSD-95 has also been studied by employing kinases capable of phosphorylating PSD-95 and subsequently identifying modified sites (Chetkovich et al., 2002, Du et al., 2009, Morabito et al., 2004), however, this approach has restricted site specificity and requires prior knowledge of the implicated kinases. We have previously introduced site-specific phosphorylations in single PDZ domains of PSD-95 by semi-synthetic strategies, and studied the effect by biochemical binding assays to a large library of C-terminal peptide interaction partners (Pedersen et al., 2017). This demonstrated how specific phosphorylations fine-tune regulation of interactions with the NMDA receptor and Stg. However, limitations of the semi-synthetic approach prevented the exploration of the effect of phosphorylation on interdomain interactions found in the full-length (FL) of PSD-95. Recently, we showed that the tandem PDZ1-2 of PSD-95 was necessary for phase separation with Stg, as single PDZ domains alone did not phase separate with Stg (Zeng et al., 2019). This illustrates the importance of interdomain interactions in the molecular mechanisms of PSD proteins.

The ability to site-specifically phosphorylate Ser residues genetically by amber codon suppression have provided the necessary means for studying phosphorylation of larger proteins (Rogerson et al., 2015, Xie and Schultz, 2005). Here, we use amber codon suppression to site-specifically phosphorylate all six Ser residues in PSD-95. We show that we can site-specifically phosphorylate multiple positions in single (PDZ1), tandem (PDZ1-2) and FL PSD-95, also revealing that some sites are more prone to amber codon suppression than others with differences found on both site and protein level. We furthermore study the effect of phosphorylation on the dynamics of the PSD, both by biochemical binding assays and by LLPS. In the latter, we show that the interdomain interactions of FL PSD-95 affect the phase separation with protein ligands. This supports the importance of studying FL proteins in contrast to single domains as the contribution of multivalent interactions is acknowledged. By combining genetically encoded site-specific phosphorylations with phase separation studies, we provided a novel way to study the effect of phosphorylation on the dynamics of the PSD.

## RESULTS

### Site-specific phosphorylation of PSD-95

To explore the feasibility of genetically introducing phosphoserine (pSer) in PSD-95, we first wanted to introduce pSer at all six Ser positions of PDZ1 in the PDZ1 single and PDZ1-2 tandem proteins of PSD-95. To do this, *E. coli* BL21Δ*serB* cells were co-transformed with the previously described single high-copy plasmid pKW2 EF-Sep (Rogerson et al., 2015) and plasmids carrying the protein coding sequences containing an amber (TAG) codon at the desired positions. The pKW2 EF-Sep plasmid contains the optimized aminoacyl tRNA synthetase and tRNA pair SepRS(2)/pSer tRNA(B4)_CUA_ specific for pSer incorporation and orthogonal to the endogenous synthetase and tRNA pairs of *E. coli*, which are responsible for incorporating the remaining natural amino acids into the proteins. The plasmid furthermore contains the elongation factor (EF)-Sep optimized for pSer incorporation (Park et al., 2011, Rogerson et al., 2015). The *E. coli* used for protein expression, BL21Δ*serB*, lack phosphoserine phosphatase preventing cleavage of the introduced phosphorylated amino acid while also maintaining an intracellular pool of pSer (Park et al., 2011). For the incorporation, 1 mM pSer was added to the media, and upon induction with 0.5 mM isopropyl β-D-1-thiogalactopyranoside (IPTG), phosphorylated variants of PDZ1 and PDZ1-2 were expressed and pSer introduced at the desired positions in response to the TAG codons.

We successfully introduced pSer at four different positions in PDZ1 and PDZ1-2 (Figure 1B,C). Ser73 and Ser78 were site-specifically phosphorylated in both PDZ1 and PDZ1-2 and the proteins purified in high yields (Figure 1B). Interestingly, two positions were only successfully phosphorylated in either one of the two protein variants, Ser142 in PDZ1 and Ser116 in PDZ1-2, respectively. PDZ1 pSer116 expressed as insoluble protein in inclusion bodies and purification resulted in low yield and purity, where a mixture of both WT and pSer mass was observed, thus the protein variant was excluded from further studies. Moreover, phosphorylation of Ser93 and Ser131 was not achieved in either of the PDZ1 pr PDZ1-2.

The positions in PDZ1 that were successfully phosphorylated in the PDZ1 and PDZ1-2 protein variants were then attempted phosphorylated in FL PSD-95 (amino acids 60-724). Here, amber codon suppression of all four positions (Ser73, Ser78, Ser116 and Ser142) resulted in expression of phosphorylated protein, albeit in lower yields (Figure 1B). FL PSD-95 was furthermore phosphorylated in position Ser295, which is located in the linker region between PDZ2 and PDZ3. The expression efficiency was very low, which resulted in a yield of less than 1 mg protein pr. liter expression culture (mg/L) (Figure 1B). Furthermore, poor efficiency of amber codon suppression at positions Ser425 and Ser561 in FL PSD-95 was observed, and the proteins were not purified (Figure 1B). Two positions in PDZ2 of the tandem PDZ1-2 were successfully suppressed and phosphorylated, Ser173 and Ser217, but not pursued further.

The yield of the phosphorylated protein variants of PDZ1, PDZ1-2 and FL PSD-95 decreased with protein size, which aligned with the observed expressions, and purification of PDZ1 WT resulted in five-fold higher yield than FL PSD-95 WT (Figure 1B). A total of 14 protein variants of PDZ1, PDZ1-2 and FL PSD-95 were generated, 11 of these phosphorylated at five different positions (Figure 1C).

### Characterization of protein variants and identification of interesting sites

The secondary structure of the proteins was evaluated by circular dichroism (CD) and the introduced phosphorylations did not affect the secondary structure (Figure 2A). Moreover, protein phosphorylation did not affect the stability of the protein variants, as measured by assessing the thermal denaturation of the proteins (**Supplementary Figure 1**).

**Figure 2:**
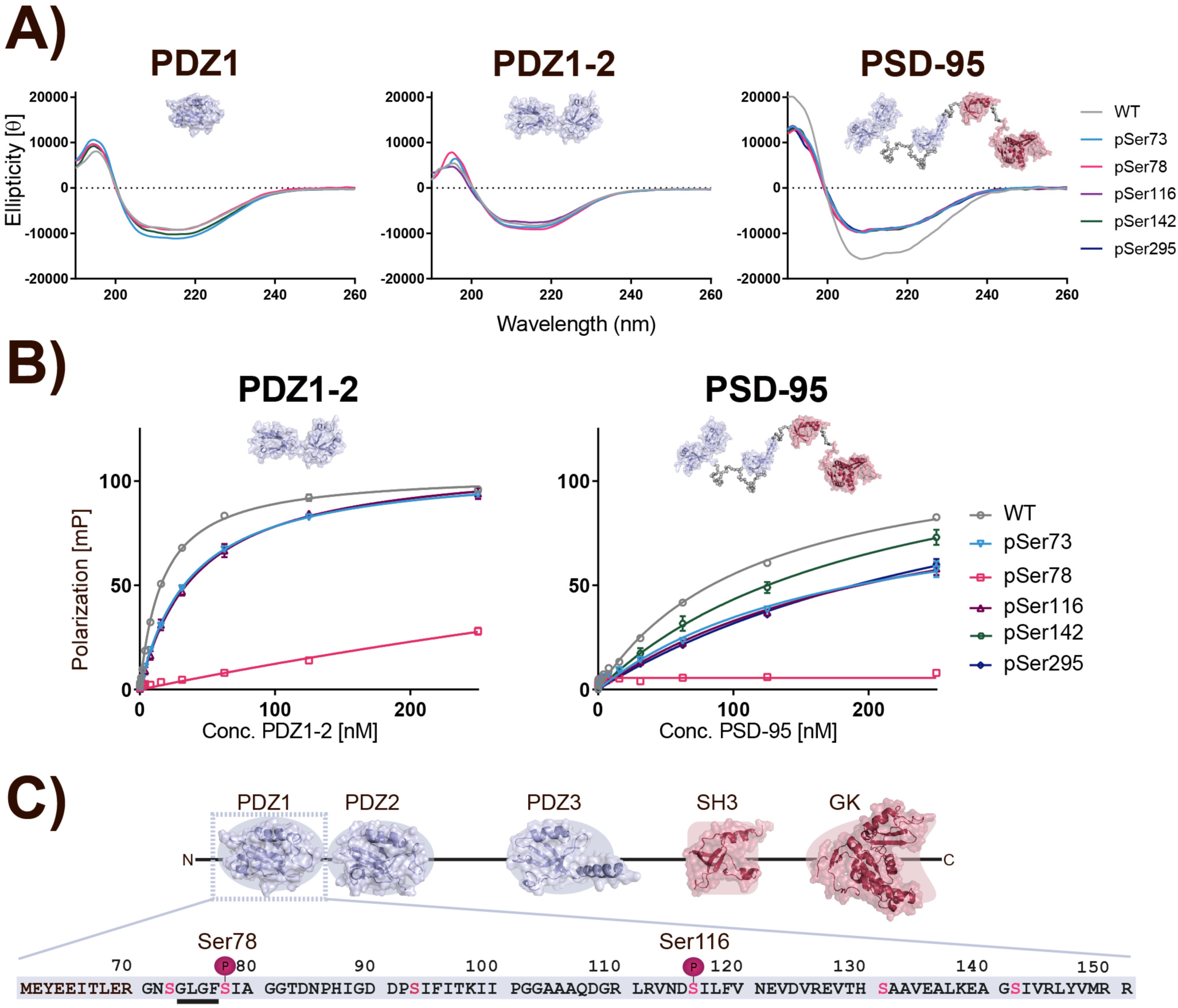
Characterization protein variants and identification of interesting sites. **A)** CD spectra for all purified protein variants of PDZ1, PDZ1-2 and PSD-95. Three accumulated spectra were measured at 25 ⁰C from 260-190 nm and the molar ellipticity calculated. **B)** Protein variants of PDZ1-2 and PSD-95 were tested for binding to peptide inhibitor AVLX-144 in a FP saturation assay. Here, 10 nM of Cy5-labeled AVLX-144 was saturated with protein in a concentration dependent manner with a maximum concentration of 250 nM protein. The milli-polarization (mP) was measured for all concentrations in triplicates, background fluorescence subtracted and the data fitted to a one-sided binding model from which *K*_d_ values were calculated. **C)** Illustration of PSD-95 and its five domains with the sequence of PDZ1 highlighted. Here, the two phosphorylation sites, Ser78 and Ser116, are illustrated and the GLGF repeat underlined.

To examine the influence of the site-specific phosphorylations for binding to known interaction partners and ligands, we measured the binding affinity of PDZ1-2 and FL PSD-95 protein variants to the fluorescently labeled dimeric peptide inhibitor AVLX-144 (Bach et al., 2012) by fluorescence polarization (FP). In a concentration dependent manner, ligand saturation with protein was measured and binding constants (*K*_d_) calculated (Figure 2B). Both protein variants bound the dimeric peptide inhibitor with nanomolar affinities (**Supplementary Table 2**). All phosphorylations were observed to decrease the binding affinity to AVLX-144 and interestingly, both protein variants phosphorylated on Ser78 did not bind to AVLX-144. Ser78 is located in the carboxylate binding site of the PDZ1 domain (Figure 2C), which can explain the dramatic effect on peptide binding. Ser116 is located in the acidic surface of the PDZ1 domain, and phosphorylation of this site also resulted in reduced binding affinity of AVLX-144.

### Identification of site-specific phosphorylations

The insertion of pSer was characterized as a mass change of 80 Dalton (Da) corresponding to the phosphate group and determined by liquid chromatography-mass spectrometry (LC-MS) (Figure 3**; Supplementary Figure 2;3;4**). The introduction of the site-specific phosphorylations was verified by LC-coupled tandem MS (LC-MS/MS). The proteins were purified to above 90 %, as determined by reverse phase ultra-performance liquid chromatography (UPLC) (**Supplementary Figure 2;3;4**), and cleaved by enzymatic digestion in order to simplify fractionation, ionization, and fragmentation. By tandem-MS the incorporation of pSer at all positions except pSer295 in FL PSD-95 was confirmed with high localization probabilities (>0.9), which is of great importance for the study of the effect of site-specific phosphorylation and validates the approach (Figure 3**; Table 3; Supplementary Figure 2;3;4**). The sites furthermore displayed high intensities and delta scores, demonstrating that they were the most confident identifications in the individual samples. Altogether, this validates the approach of genetic incorporation of pSer intro PSD-95.

**Figure 3:**
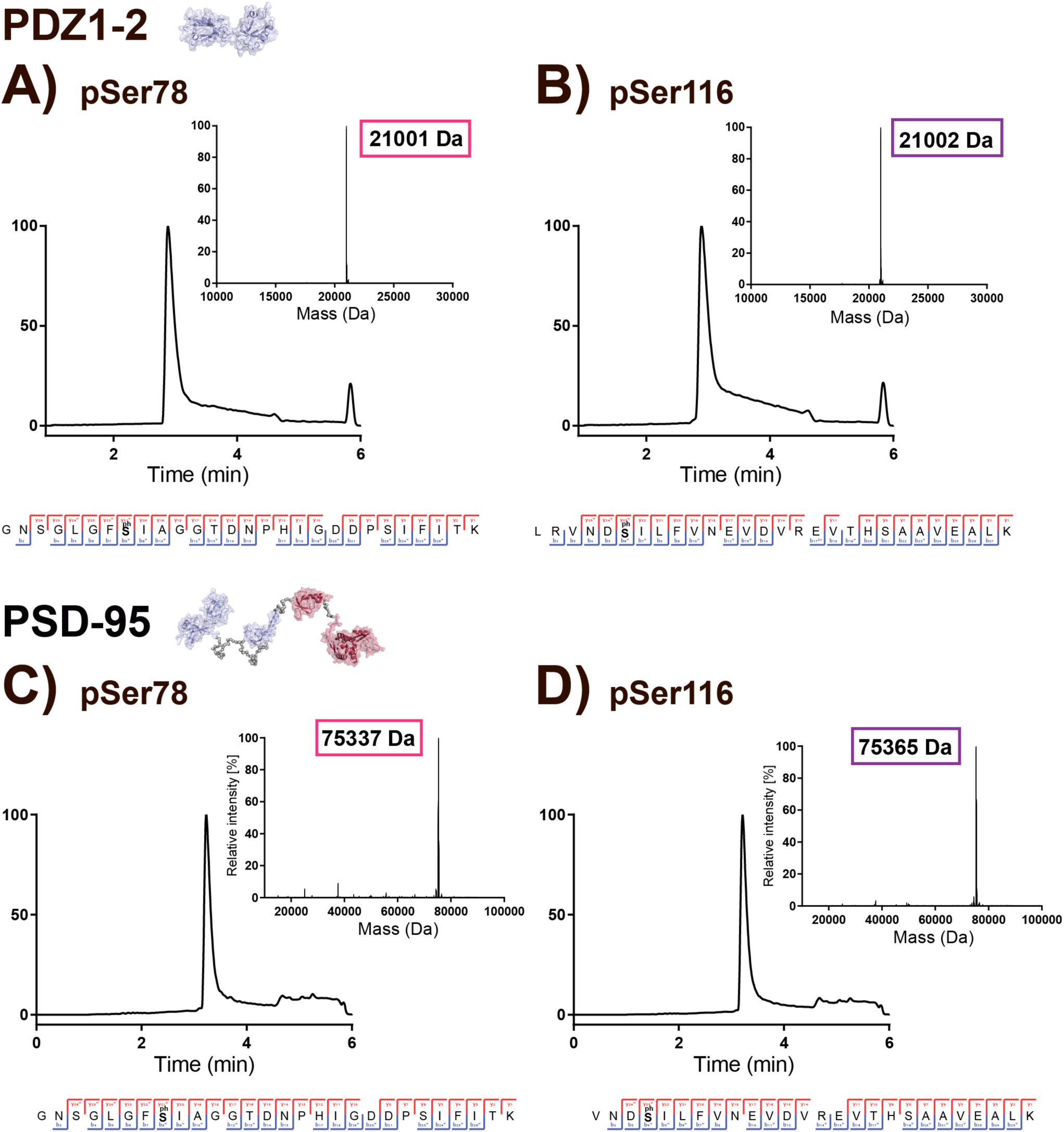
Identification of site-specific phosphorylations. **A)** PDZ1-2 pSer78 protein variant. LC-MS spectrum and mass derived from the deconvoluted spectrum of the purified PDZ1-2 pSer78 protein variant. The observed pSer mass is highlighted in color. The expected mass for the PDZ1-2 pSer protein variant was 21004 Da. Below the spectra is the identified phosphorylated peptide from the LC-MS/MS analysis of the purified protein. The annotated spectrum is found in **Supplementary Figure 3**. **B)** PDZ1-2 pSer116 protein variant. The annotated spectrum is found in **Supplementary Figure 3**. **C)** PSD-95 pSer78 protein variant. LC-MS spectrum and mass derived from the deconvoluted spectrum of the purified PSD-95 pSer78 protein variant. The observed pSer mass is highlighted in color. The expected mass for the PSd-95 pSer protein variant was 75349 Da. Below the spectra is the identified phosphorylated peptide from the LC-MS/MS analysis of the purified protein. The annotated spectrum is found in **Supplementary Figure 4**. **D)** PSD-95 pSer116 protein variant. The annotated spectrum is found in **Supplementary Figure 4**.

### Effect of phosphorylation of Ser78 and Ser116 on binding to stargazin and GluN2B

To further evaluate the effect of phosphorylation, all pSer variants were analyzed for the binding to ligands by isothermal titration calorimetry (ITC) and FP assays. Through its interactions with the C-terminal of subunits of the NMDA receptor and Stg, the auxiliary subunit of the AMPA receptor, PSD-95 regulates the two most prevalent ionotropic glutamate receptors (iGluRs) at the PSD and these interactions were the primary interactions of interest.

The effect of phosphorylation on the binding to Stg was measured by ITC for all protein variants. Here, the PDZ-binding motif (PBM) and C-terminal of Stg (amino acids 203D-323V) was employed as the protein ligand, which all proteins were observed to bind with micromolar affinities (Figure 4A,B**; Supplementary Figure 5)**. The binding affinity increased with protein size and FL PSD-95 WT was observed to bind Stg with an affinity of 1.2 µM. This was almost a 100-fold higher than the observed 100 µM binding affinity of PDZ1 WT to Stg (Figure 4B**; Supplementary Figure 5A**) and 10-fold higher than PDZ1-2 WT, which bound with an affinity of 12 µM (Figure 4A,B). PDZ1 WT bound Stg with very low affinity and the variant phosphorylated on Ser78 was observed not to bind (**Supplementary Figure 5A)**. However, phosphorylation of Ser78 in PDZ1-2 and FL PSD-95 did not result in this change of binding affinity (Figure 4A,B), which can be explained by an earlier finding that the Stg tail does not bind to the canonical target recognition groove of PDZ1 in the Stg/PDZ1-2 or Stg/FL PSD-95 complexes(Zeng et al., 2019). Phosphorylation of Ser116 in both PDZ1-2 and FL PSD-95 slightly enhanced binding to Stg, seen as change in the binding slope and thereby had the opposite effect as seen for phosphorylation of Ser78 in PDZ1 (Figure 4A,B**; Supplementary Figure 5A**). Phosphorylation of Ser73 and Ser142 in PDZ1 enhanced the weak binding to Stg (**Supplementary Figure 5A**), and phosphorylation of Ser73 in PDZ1-2 negatively affected binding to Stg (**Supplementary Figure 5B**). Phosphorylation of Ser73 and Ser142 in FL PSD-95 did not alter the binding affinity to Stg (**Supplementary Figure 5C**).

**Figure 4:**
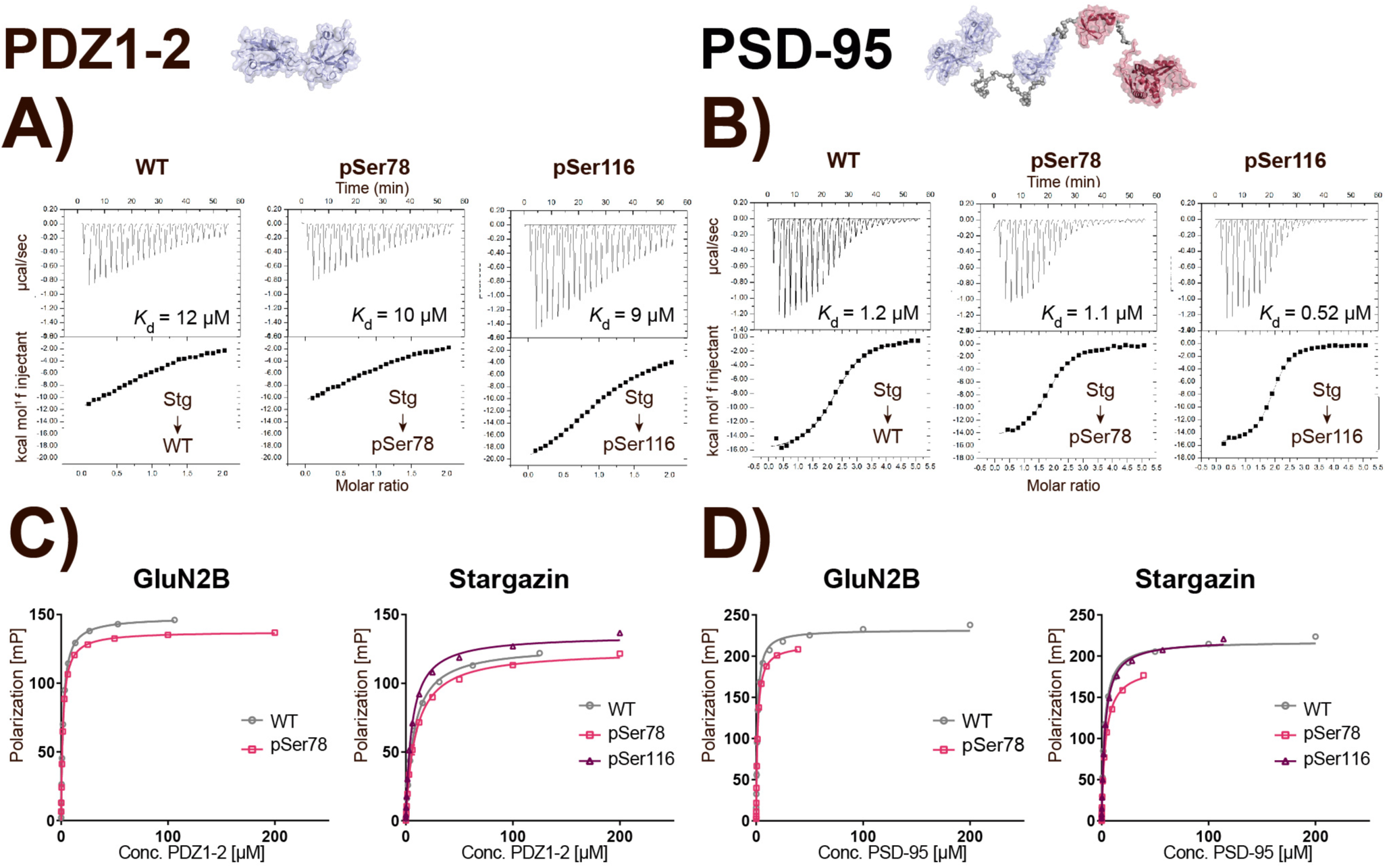
Effect of phosphorylation of Ser78 and Ser116 on binding to stargazin and GluN2B. **D)** ITC-based measurements of the PBM and C-terminal of stargazin (Stg) binding to PDZ1-2 WT, pSer78 and pSer116. 250 µM Stg was titrated into 25 µM PDZ1-2 in the cell and binding constants (*K*_d_) were calculated. **E)** ITC-based measurements of the PBM and C-terminal of stargazin (Stg) binding to PSD-95 WT, pSer78 and pSer116. 250 µM Stg was titrated into 10 µM PSD-95 in the cell and binding constants (*K*_d_) were calculated. **F)** PDZ1-2 binding to GluN2B and stargazin peptide ligands measured in a saturation FP assay. Here, 200 nM of the TAMRA-labeled C-terminal peptide ligand was saturated with protein in a concentration dependent manner with a maximum concentration of 200 µM protein, if possible. The mP was measured for all concentrations in triplicates, background fluorescence subtracted and the data fitted a to a one-sided binding model from which the *K*_d_ values were calculated. The *K*_d_ values are listed in supplementary table 5 and highlighted Ser78 (*) and Ser116 (*). **G)** PSD-95 binding to GluN2B and stargazin peptide ligands measured in a saturation FP assay. The conditions described in C) were used.

The effect on ligand binding was also measured by FP using C-terminal TAMRA-labeled peptide ligands, representing protein interaction partners (Pedersen et al., 2017). A total of five peptide ligands were employed and tested for binding to specific phosphorylated protein variants (**Supplementary Table 3**). All proteins bound the ligands with micromolar affinities, and the affinity increased with protein size (**Supplementary Table 4**), as was also observed by ITC. PDZ1 WT was observed to bind GluN2B with an affinity of *K*_d_ = 12.4 µM, however, phosphorylation of Ser78 disrupted this binding (**Supplementary Figure 6A, top middle**). Likewise, PDZ1 WT bound to Stg peptide with an affinity of 27.3 µM, but phosphorylation of Ser78 disrupted this binding (**Supplementary Figure 6A, bottom middle**). Phosphorylation of Ser78 in PDZ1-2 and FL PSD-95 did however not display the same dramatic effect on binding to both GluN2B and Stg peptides (Figure 4C,D). PDZ1-2 and FL PSD-95 WT proteins bound GluN2B with almost equally high affinities of 1.9 µM and 1.2 µM, respectively, but phosphorylation of Ser78 in both proteins did not affect this binding (Figure 4C,D **left graphs; Supplementary Table 4**). Both PDZ1-2 and FL PSD-95 WT were also observed to bind Stg with high affinities of 7.5 µM and 2.7 µM, respectively, and here, phosphorylation of Ser78 negatively affected the binding, however not significantly (Figure 4C,D **right graphs; Supplementary Table 4**). Phosphorylation of Ser116 in PDZ1-2 was observed to slightly enhance binding to Stg by ITC, and this was also observed in the FP assay (Figure 4C**, right graph; Supplementary Table 4**). Phosphorylation of Ser116 in FL PSD-95 slightly decreased binding to Stg, albeit not significantly (Figure 4D**, right graph; Supplementary Table 4**).

Phosphorylation of Ser73 in PDZ1 negatively affected binding to both GluN2A and GluN2B subunits of the NMDA receptor (**Supplementary Figure 6A, top and bottom left**), as previously observed (Pedersen et al., 2017). The effect was not as significant for phosphorylation of the same site in the tandem or FL protein variants (**Supplementary Figure 6B,C (top and bottom left)**). Phosphorylation of Ser142 in PDZ1 and FL PSD-95 did not affect binding to the C-terminal peptides representing the Kv1.4 or Kv1.7 potassium channels (**Supplementary Figure 6A, right top and bottom**).

Interestingly, increased binding affinities of ligands were observed with an increase in protein size, which could be a result of multiple factors including the multivalent binding nature of the FL protein in contrast to particularly the single PDZ domain. The PDZ3-SH3-GK module of FL PSD-95 functions as another binding site for ligands and additionally FL PSD-95 can experience target binding-induced multimerization in contrast to single PDZ domains, which could increase the avidity of the binding. Altogether this could lead to weaker phenotypes for the phosphorylated FL PSD-95 protein variants relative to the single domain variants, observed as a less impact of phosphorylation on measured binding affinities.

### Effect of phosphorylation on phase separation with GluN2B and stargazin

We have previously shown that the tandem domain of PSD-95 is necessary for the phase separation with Stg, as the single domains alone did not phase separate (Zeng et al., 2019). To test if the introduced phosphorylations in PDZ1-2 and FL PSD-95 affected the phase separation with both GluN2B and Stg, sedimentation-based experiments were used to assess the formation of condensates. The protein ligands were mixed individually with the phosphorylated protein variants and the ratio of protein in the aqueous (supernatant) and condensed (pellet) phases analyzed by gel electrophoresis.

Both PDZ1-2 and FL PSD-95 were observed to form condensates via LLPS with GluN2B at 10 and 5 µM, respectively (Figure 5A **top; Supplementary Figure 6D**). Phosphorylation of Ser78 in PDZ1-2 was observed to inhibit phase separation with GluN2B as less protein was seen in the condensate in comparison to phase separation with the WT protein variant. Interestingly, measured by FP phosphorylation of Ser78 in PDZ1-2 did not affect binding to GluN2B (Figure 4C) but the effect on phase separation was clearly noticeable. The same inhibitory effect was observed by ITC and FP for PDZ1 pSer78, but as single domains do not phase separate, this was not evaluated. Neither phosphorylation of Ser116 in PDZ1-2 nor phosphorylation of Ser73, Ser142 and Ser295 in FL PSD-95 had an effect on phase separation with GluN2B (**Supplementary Figure 6D**). Phase separation of both PDZ1-2 and FL PSD-95 with Stg was also observed (Figure 5A **(bottom),B; Supplementary Figure 6E**) and phosphorylation of Ser78 in PDZ1-2 clearly inhibited the formation of condensates with Stg. The effect was less significant for FL PSD-95 pSer78, but more FL PSD-95 pSer78 and Stg was observed in the aqueous phase, indicating inhibition of condensate formation. On the contrary, phosphorylation of Ser116 in PDZ1-2 promoted phase separation with Stg, and this effect was also observed for FL PSD-95 pSer116, however again to a lesser extent. None of the remaining sites resulted in altered formation of condensates via LLPS.

**Figure 5:**
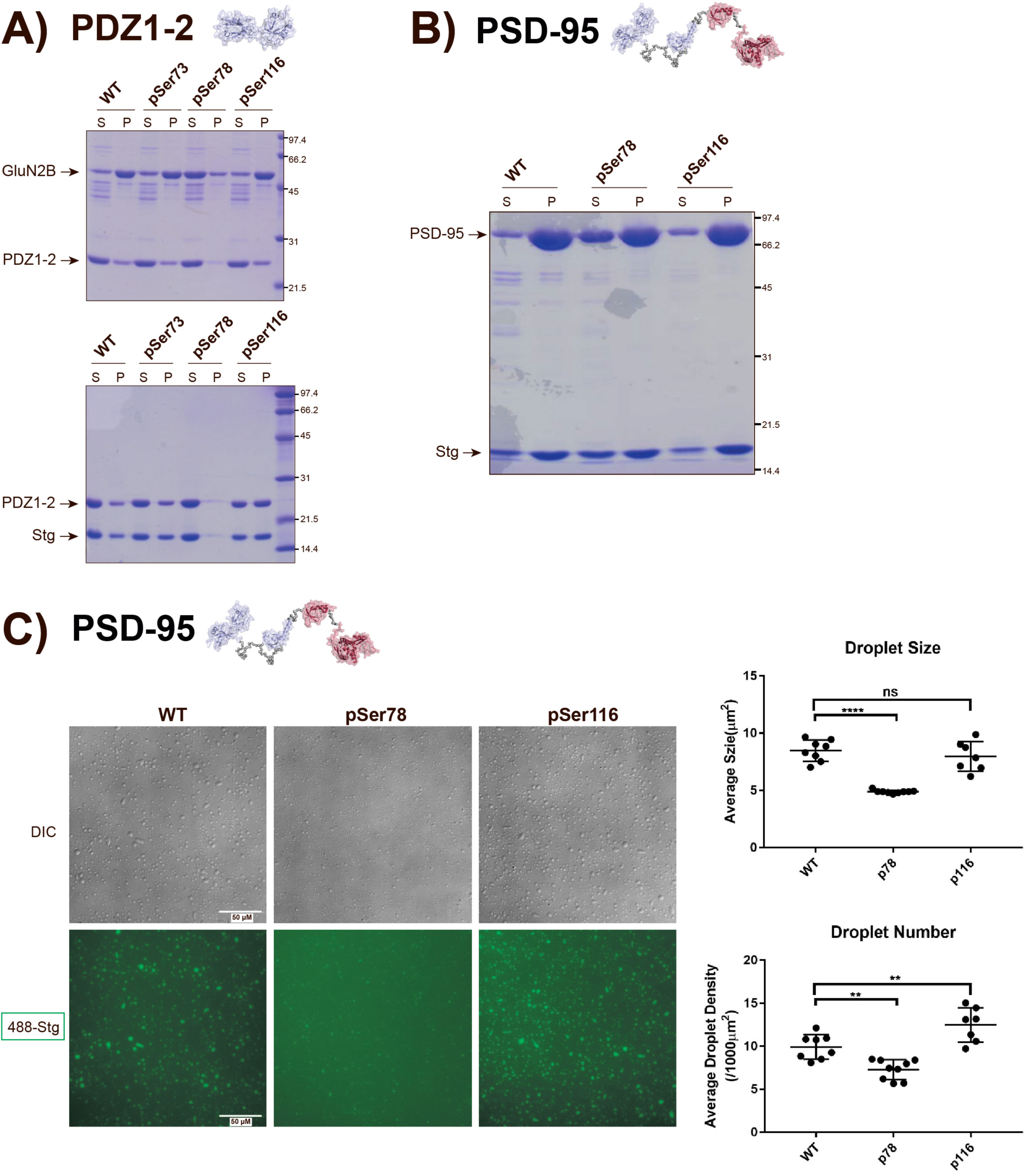
Effect of phosphorylation on phase separation with GluN2B and stargazin. **A)** PDZ1-2 (10 µM) was mixed with GluN2B protein (10 µM) (top) and stargazin protein (Stg) (bottom), equilibrium reached and the mixture spun down. Hereafter, the supernatant (S) and resuspended pellet (P) was loaded on a gel, which was coomassie stained to visualize the protein bands. Bands corresponding to PDZ1-2, GluN2B and Stg are indicated by an arrow. For sedimentation assays with PDZ1-2, 10 µM PDZ1-2 and 10 µM GluN2B was used, and 75 µM protein and 75 µM Stg. **B)** PSD-95 was mixed with stargazin protein (Stg), equilibrium reached and the mixture spun down. Hereafter, the supernatant (S) and resuspended pellet (P) was loaded on a gel, which was coomassie stained to visualize the protein bands. Bands corresponding to PSD-95 and stargazin are indicated by an arrow. For sedimentation assays with PSD-95, 5 µM pSer78 and pSer116 protein and 12.5 µM stargazin was used. **C)** DIC (top) and fluorescence (bottom) images showing the phase separation of PSD-95 WT, pSer78 and pSer116 (5 µM) with Stg (12.5 µM). For the fluorescence images Stg was labeled with Alexa Fluor 488. The scale of 50 µM is indicated.

We also observed LLPS of FL PSD-95 and Stg under light microscopy (Figure 5C). Differential interference contrast (DIC) images revealed numerous spherical-shaped and micron-sized droplets of FL PSD-95 WT and Stg, and fluorescence images showed that the droplets were enriched with Stg. Ser78 phosphorylation in FL PSD-95 was observed to inhibit the formation of lipid-droplets, as also seen by sedimentation assays. Phosphorylation of Ser116 was in contrast observed to promote concentration of protein in the condensed phase, which supports the sedimentation assay.

## DISCUSSION

PSD-95 is one of the most abundant proteins of the PSD, and orchestrates signaling of particularly the AMPA and NMDA receptors through interactions fine-tuned by PTMs, such as phosphorylation. The interactions are primarily mediated by the three PDZ domains of PSD-95, and due to the multivalent binding mode of PSD-95, the interactions are highly complex. The effect of site-specific phosphorylation on the complex interactions are however largely unknown and here we aimed to evaluate this by combining biochemical binding assays with phase separation studies of genetically phosphorylated protein variants of PSD-95.

Here, we generated 11 site-specifically Ser phosphorylated protein variants of PSD-95 using amber codon suppression and found that the introduced phosphorylations did not affect the protein fold (Figure 1, Figure 2A). We have previously site-specifically phosphorylated single PDZ domains of PSD-95 using a semi-synthetic approach (Pedersen et al., 2017) but here Ser residues in both the single, tandem and FL PSD-95 were phosphorylated in both high yields and purity. This indicated the strength of the approach for generation of many phosphorylated protein variants. With amber codon suppression, theoretically all Ser residues can be phosphorylated (Rogerson et al., 2015), but we observed differential suppression depending on the site of introduction (Figure 1B). Two positions in PDZ1 of the single and tandem protein variants were not successfully suppressed (Ser93 and Ser131) and furthermore, two positions were only successfully phosphorylated in either one of the variants (Ser116 and Ser142). Interestingly, all attempted positions phosphorylated in PDZ1 of FL PSD-95 were however successfully suppressed. Site-dependable suppression of amber codons is a common challenge in the field of genetic incorporation of non-natural amino acids, and can be the result of various factors(Chin, 2017). The sequence context of the inserted amber codon can affect the suppression efficiency with nucleotides both upstream and downstream of the amber codon potentially limiting the suppression efficiency (Schwark et al., 2018, Xu et al., 2016). The localization of the amber codon within a protein as well as the stability of the expressed protein can furthermore affect the expression of non-canonical amino acid containing proteins, affecting the protein yield (Zhu et al., 2019). In contrast to the natural mechanism for post-translational modification of a protein, expression of phosphorylated proteins using amber codon suppression requires the protein to fold with the inserted phosphorylation, which potentially could affect the expression efficiency. These challenges can be addressed by exploring the usage of other bacterial strains, where e.g. amber stop codons have been recoded and release factor (RF) proteins have been deleted (Heinemann et al., 2012, Lajoie et al., 2013, Mukai et al., 2015), to increase the expression of phosphorylated protein. Many of these strains have however suffered from low fidelity, with a high insertion of canonical amino acids in response to the amber codon, slow growth rates and an unhealthy metabolism. Various systems have furthermore been developed for pSer insertion in both proteins (Lee et al., 2013, Pirman et al., 2015) and peptides (Barber et al., 2018), and could be evaluated to determine the cause for the differential site suppression seen here. This has been done recently using a further developed pSer system in a healthy RF-deficient strain (Zhu et al., 2019). As we only saw the differential site-dependence for the single and tandem domains, the differential suppression could however indicate that other factors such as stability or functionality of the modified protein affected the outcome.

All phosphorylated proteins were purified in high purity, albeit the yields decreased with increased protein size (Figure 1B). The genetic insertion of pSer is a means of competition between the natural translation machinery, reading the amber codon as a stop codon, and the orthogonal, reading the amber codon for pSer insertion, and the expression of truncated protein product is therefore evident as a factor affecting the overall protein yield (Chin, 2017). Expression of truncated product was seen for all sites attempted suppressed in the FL PSD-95 protein variant and the expression of truncated protein products were greater for positions 295, 425 and 561. These were therefore either purified in very low yield or not at all. The truncated products of the single and tandem domains were undetectable due to their small size not visible by SDS-PAGE, and the expression of truncated product could therefore not be evaluated for these protein variants. It is hence unclear if the lower yield of FL PSD-95 protein in contrast to the single and tandem domains were primarily due to loss of protein during purification or also differential expression efficiency with lower expression of FL PSD-95 phosphorylated protein. The site-specific phosphorylations were verified using tandem-MS with all sites showing both high localization probabilities and intensities (Figure 3; Table 1**; Supplementary Figure 2;3;4**). The results highly correlated with the expression and purification results, as the FL PSD-95 pSer295 protein variant, which was insufficiently suppressed, was more difficult to detect by tandem-MS. This suggested that better and more accurate phosphorylation was achieved with high suppression efficiency.

**Table 1:** LC-MS/MS derived data values for all protein variants. For each protein variants of PDZ1, PDZ1-2 and PSD-95, the localization probability, intensity, score and delta score is listed.

| Protein | Site | Localization probability | Intensity | Score | Delta score |
| --- | --- | --- | --- | --- | --- |
| **PDZ1** | pSer73 | 0.9855 | 6.4E+10 | 186.58 | 175.94 |
|  | pSer78 | 0.9922 | 9.01E+10 | 198.20 | 187.23 |
|  | pSer142 | 0.9998 | 2.15E+11 | 197.37 | 180.31 |
| **PDZ1-2** | pSer73 | 0.9925 | 1.5E+10 | 235.17 | 224.16 |
|  | pSer78 | 0.9814 | 1.79E+10 | 210.35 | 199.28 |
|  | pSer116 | 1 | 6.38E+09 | 129.90 | 117.13 |
| **PSD-95** | pSer73 | 0.9885 | 1.18E+10 | 135.40 | 125.94 |
|  | pSer78 | 0.9988 | 1.31E+10 | 160.90 | 149.38 |
|  | pSer116 | 0.9993 | 3E+10 | 150.12 | 136.20 |
|  | pSer142 | 1 | 2.89E+10 | 116.28 | 100.75 |
|  | pSer295 | NA | NA | NA | NA |

When evaluating the effect of the site-specific phosphorylations by ITC and FP assays, as well as phase separation, a clear tendency was seen (Figure 4; Figure 5). Here we observed an increased binding affinity for both peptide and protein ligands with an increase in protein size, with highest affinity for FL PSD-95, then PDZ1-2 and lowest for PDZ1, albeit this was accompanied by a weaker phenotype of the introduced phosphorylations. FL PSD-95 protein variants were found to bind ligands with up to 10-fold higher affinity than PDZ1 variants, but the effect of phosphorylation of the multivalent FL PSD-95 was less pronounced than phosphorylation of the single PDZ1 domain. Phosphorylation of Ser78 in the PDZ1 protein variant resulted in no binding of Stg or GluN2B ligands. Observed by biochemical binding assays, this effect was less significant for phosphorylation of the same site in both the PDZ1-2 and FL PSD-95 protein variants. The effect was more pronounced when observed by phase separation, especially for PDZ1-2, where the results also strongly indicated that phosphorylation of Ser116 promoted phase separation with Stg only, functionally contrary to phosphorylation of Ser78.

The increased binding affinity and less significant effect of phosphorylation of FL PSD-95 in contrast to PDZ1 could be attributed to compensatory mechanisms of the additional protein domains as well as the multivalent binding mode of FL PSD-95 (Zeng et al., 2019). PSD-95 can simultaneously bind multiple ligands as it contains three PDZ domains, and this likely contributed to the increased binding affinity that we observed. This suggests that the multiple ligand binding sites resulted in the introduced phosphorylations having a minor effect on the interaction with the various binding partners, but were however also driving forces for the formation of LLPS condensates. The ability of proteins to phase separate and the threshold for phase separation relies on multiple factors including the domain valency and intrinsically disordered regions in the protein sequence (Li et al., 2020, Chen et al., 2020, Banani et al., 2017). Multivalency drives LLPS and increased network valency tends to enhance phase separation (Li et al., 2012). Despite clear formation of condensates, the lower threshold for phase separation of PDZ1-2 protein variants with the two ligands could therefore explain the very visible effect of phosphorylation. Likewise, the added valency from the additional domains in FL PSD-95 could explain lack of a clear observable effect of phosphorylation of the multidomain protein.

Previous studies have shown that Stg uses its entire C-terminal (amino acids 203-323) incl. the PBM to bind to PSD-95 via specific and multivalent interactions, which also governs the phase separation with PSD-95 (Zeng et al., 2019). Namely, the disordered Arg-rich C-terminal has been found to bind PDZ1 and the PBM to bind PDZ2 of PSD-95. Furthermore, Stg binds to PDZ3 of PSD-95. This multivalent binding mode of Stg to PSD-95 could also explain the less visible effect of Ser78 phosphorylation of both the PDZ1-2 tandem domain and FL PSD-95 in contrast to phosphorylation of the same position in PDZ1, as a second binding site of Stg to FL PSD-95 would contribute with an increased binding affinity and shield the effect of the introduced phosphorylation. This would explain the dramatic effect of phosphorylation of Ser78 in PDZ1, but could be further investigated. Likewise, the interaction between PDZ1 and the Arg-rich C-terminal of Stg is governed by the negatively charged surface of PDZ1 (Zeng et al., 2019), and as Ser116 is near this surface, phosphorylation would increase the negative charges and promote interaction.

Interestingly, our results identify two novel phosphorylation sites in PDZ1 of PSD-95, namely Ser78 and Ser116, with opposing effects on ligand binding and phase separation. Phosphorylation of Ser78 inhibited binding of both GluN2B and Stg, whereas phosphorylation of Ser116 promoted binding with Stg only, which could indicate importance in receptor distribution, removal and accumulation, and thereby synaptic transmission. The functionality of these phosphorylations could be further investigated by introducing non-binding mutations or genetically encoding a non-hydrolysable analogue for cellular studies.

Overall, the data presented proved the feasibility of amber codon suppression for genetic introduction of site-specific phosphorylations in three different protein variants of PSD-95, and in combination with both biochemical assays and phase separation was found to be a good approach for the identification of novel phosphorylations sites.

## ACKNOWLEDGEMENTS

The work was supported by grants from the SUND Graduate School and The Center for Biopharmaceuticals to KS and grants from Research Grant Council of Hong Kong (AoE-M09-12 and C6004-17G) to MZ. MZ is a Kerry Holdings Professor of Science and a Croucher Foundation Senior Fellow. The construct pKW2 EF-Sep was kindly provided by Professor Jason Chin, Cambridge, UK. Nikolaj Riis Christensen and Mette Ishøy Rosenbaum contributed with scientific guidance and discussion.

## AUTHOR CONTRIBUTIONS

MVP, TLJ, and SB expressed and purified the pSer protein variants, and characterized the proteins. MVP and SB performed all FP experiments. MVP and SCBL performed LC-MS/MS analysis. CM purified protein for LLPS and XC performed ITC and LLPS experiments. MVP, XC and SCBL analyzed all the data. MVP, XC, LSC, MZ and KS designed the research. MVP and KS drafted the paper. All authors commented on the paper.

## STAR METHODS

### Plasmid generation

Three different variants of the human PSD-95 protein were used in this study, PSD-95 61-724, PSD-95 PDZ1 and PSD-95 PDZ1-2. All proteins were expressed with a C-terminal poly-histidine tag (6xHis) from a pCDF-1b vector (Novagen) with spectinomycin resistance and under the control of a T7 lac promoter. The DNA constructs were verified by DNA sequencing (Eurofins Genomics).

To generate pCDF-1b Trx HRV 3C PSD-95 61-724 6xHis, termed PSD-95, the protein coding sequence of PSD-95 61-724 6xHis flanked N-terminally by an overhang, *NdeI* restriction site and a HRV 3C cleavage site, and C-terminally by an *AvrII* restriction site and overhang, was purchased (ThermoFisher Scientific). The recipient vector-construct, pCDF-1b Trx *NdeI* HRV 3C PSD-95 1-724 6xHis *AvrII*, which was available in the laboratory, and the purchased sequence were digested with *NdeI* and *AvrII* restriction enzymes overnight (ON) at 37 ⁰C. The recipient vector and protein coding sequence were purified using a NucleoSpin® Gel and PCR Clean-up kit (Macherey-Nagel) and ligated together with a Rapid DNA ligation kit (Roche) in a ratio of 50 ng vector to 150 ng DNA to yield pCDF-1b Trx HRV 3C PSD-95 61-724 6xHis.

The pCDF-1b PSD-95 PDZ1 6xHis construct was generated from the previously described pRSET vector containing the recombinant fragment of PSD-95 PDZ1 flanked N-terminally by a 6xHis (Pedersen et al., 2017). By polymerase chain reaction (PCR) amplification, the N-terminal 6xHis tag was removed and replaced with an *NcoI* restriction site with primer MVM_034 (**Supplementary Table 1**), and a C-terminal 6xHis tag followed by an *AvrII* restriction site was inserted with primer MVM_035 (**Supplementary Table 1**). The PCR product was purified using a NucleoSpin® Gel and PCR Clean-up kit (Macherey-Nagel). The purified DNA and the recipient vector pCDF-1b Trx HRV 3C PSD-95 61-724 6xHis were cleaved during enzymatic digest with *NcoI* and *AvrII* ON at 37 ⁰C, purified and ligated together to generate pCDF-1b PSD-95 PDZ1 6xHis.

To generate pCDF-1b PSD-95 PDZ1-2 6xHis, the protein coding sequence of PSD-95 PDZ1-2 6xHis flanked N-terminally by an *NcoI* site and C-terminally by an *AvrII* site was purchased. The DNA and the recipient vector pCDF-1b Trx HRV 3C PSD-95 61-724 6xHis were cleaved during enzymatic digest with *NcoI* and *AvrII* ON at 37 ⁰C, purified and ligated together to generate pCDF-1b PSD-95 PDZ1-2 6xHis.

Generation of site-mutated serine residues in all constructs were conducted by quick-change PCR followed by *DpnI* digest at 37°C for 1.5 hours (hrs) (New England Biolabs). Forward and reverse primers were designed for each mutated serine position to incorporate a TAG codon instead (**Supplementary Table 1**).

Constructs containing the protein coding sequences of stargazin and GluN2B for phase separation studies were generated using standard PCR-based methods, cloned into vector containing an N-terminal Trx-6xHis-to stargazin and GB1-6xHis-to GluN2B followed by an HRV 3C cutting site. Constructs were confirmed by DNA sequencing.

### Protein expression incl. double transformation

For the overexpression of all three wild-type (WT) protein variants, *E.coli* BL21 competent cells (DE3, pLysS, Invitrogen) were transformed with pCDF-1b Trx HRV 3C PSD-95 61-724 6xHis, pCDF-1b PSD-95 PDZ1 6xHis and pCDF-1b PSD-95 PDZ1-2 6xHis, respectively, and grown on spectinomycin (100 µg/mL) containing Luria-Bertani (LB) agar-plates ON at 37 ⁰C. Single colonies were used to inoculate 100 mL LB media containing antibiotic and pre-cultures were grown ON at 30 ⁰C shaking at 210 rpm. Pre-heated LB-media containing antibiotic was inoculated with the overnight cultures to reach an optical density (OD) at 600 nm of 0.1, and expression cultures were grown at 37 °C to an OD of 0.4-0.6 before induced with a final concentration of 0.1 mM Isopropyl β-D-1-thiogalactopyranoside (IPTG). PSD-95 was overexpressed for 4 hrs at 30 °C, PSD-95 PDZ1 for 4 hrs at 37 °C, and PSD-95 PDZ1-2 ON at 18 °C. Cells were harvested by centrifugation at 10,000xG, 4 °C, 10 min (Sorvall Lynx 6000, Thermo Scientific) and the pellets were stored at −20 °C until further use.

For the overexpression of phosphoserine (pSer) variants of all three proteins, *E.coli* BL21Δ*SerB* cells (Addgene #34929) were co-transformed with serine-mutated DNA and the high-copy number pKW2 EF-Sep plasmid kindly provided by Professor Jason Chin, Cambridge, and previously described in (Rogerson et al., 2015). Double transformed bacteria were grown on LB agar-plates containing both spectinomycin (100 µg/mL) and chloramphenicol (25 µg/mL). Overexpression was carried out similarly to overexpression of WT protein, but protein expression was induced with a final concentration of 0.5 mM IPTG and media supplemented O-phospho-L-serine (Sigma) to a final concentration of 1 mM.

*E.coli* BL21 cells transformed with mouse Stargazin_CT (aa 203-323) fused to the C-terminal end of the Trx-His6-tag or rat GluN2B cytoplasmic tail (aa 1170-1482) fused to the C-terminal end of the GB1-His6-tag were cultured in LB medium at 37 ⁰C till OD600 reached 0.8-1.0. IPTG at 0.25 mM final concentration was added to induce protein expression. Stargazin_CT was expressed at 37 ⁰C for 2-3 hours, and GluN2B fragment was expressed at 16 ⁰C for 16-20 hours.

### Protein purification

Protein pellets were lysed in ice-cold lysis buffer, either B-PER™ Bacterial Protein Extraction Reagent (ThermoFisher Scientific) or 50 mM sodium phosphate (NaPi), 10 mM MgCl_2_, pH 7.4, both supplemented 25 µg/mL DNase and cOmplete protase inhibitor tablets (Roche). If lysed in 50 mM NaPi buffer, lysate was processed in a cell-disrupter apparatus (Constant System Ltd) at 26 kPsi. Hereafter, lysate was spun down by centrifugation at 30,000xG, 4 °C, 30 min, and the pH of the supernatant adjusted to 7.5 prior to filtration through a 0.45 µM filter. Protein solutions were loaded on pre-equilibrated HisTrap columns (GE Healthcare) using either a peristaltic pump or an AKTA Start system (GE Healthcare), and eluted with imidazole either isocradically using a peristaltic pump or with a gradient on an AKTA Explorer 100 Air (GE Healthcare) or AKTA Start System (GE Healthcare).

For the purification of PSD-95 6xHis, affinity purification was carried out in 25 mM HEPES, 150 mM NaCl, pH 8, 2 mM BME supplemented either 25 mM imidazole for binding or 250 mM for elution. The purest fractions were pooled and imidazole was removed by dialysis at 4 °C ON into 25 mM HEPES, 150 mM NaCl, pH 8, 2 mM BME. Hereafter, the Trx solubility tag was cleaved off with HRV 3C protease (ThermoScientific) during dialysis at 4 °C ON into 25 mM HEPES, 10 mM NaCl, pH 8, 2 mM BME before impurities were removed by anion exchange on a MonoQ column, yielding PSD-95 6xHis. For the purification of PSD-95 pSer561, an additional step of size-exclusion chromatography (SEC) (HiLoad Superdex 16/600 200 pg column) was carried out in 25 mM HEPES, 500 mM NaCl, pH 7.4.

For the purification of PSD-95 PDZ1 6xHis and PSD-95 PDZ1-2 6xHis, affinity purification was carried out in 50 mM Tris, 150 mM NaCl, pH 7.4, 2 mM BME supplemented either 25 mM imidazole for binding or 250 mM for elution. The purest fractions were pooled and impurities removed by size-exclusion chromatography (HiLoad Superdex 16/600 75 pg column) in 50 mM Tris, 150 mM NaCl, pH 7.4, 2 mM BME, yielding PSD-95 PDZ1 6xHis and PSD-95 PDZ1-2 6xHis.

Before further experiments, purified protein was dialyzed ON at 4 °C into 25 mM HEPES, 150 mM NaCl, pH 8, 2 mM BME.

Purification was followed by validating eluted fractions by sodium dodecyl sulfate polyacrylamide gel electrophoresis (SDS-PAGE) and liquid chromatography-mass spectrometry (LC-MS) (see section for protein validation), and the purest fractions were pooled and up-concentrated in Amicon-Ultra centrifugal filters (Sigma-Aldrich).

The final purified protein variants were evaluated by LC-MS and reverse phase ultra-performance liquid chromatography (RP-UPLC) (see section for protein validation). The protein concentration was determined using absorbance at λ280nm measured on a NanoDrop 1000 spectrometer (Thermo Fisher Scientific). The protein yield was calculated based on protein concentration (mg/mL), the end-volume of purified protein, the purity and the volume of expressed cell culture, resulting in a protein yield in mg/L culture.

Recombinant Trx-His6-Stargazin_CT were freshly purified using a nickel-NTA agarose affinity column followed by a size-exclusion chromatography (Superdex 75 from the GE Healthcare) with a column buffer containing 50 mM Tris, pH 7.8, 300 mM NaCl, 1 mM EDTA, 1 mM DTT. The N-terminal Trx-His6-tag was cleaved by HRV 3C protease and removed by an additional step of ion exchange chromatography (Recourse S, GE Healthcare). Finally, a desalting column was used to exchange the protein into the buffer containing 50 mM Tris pH 7.8, 100 mM NaCl, 1 mM EDTA, 1 mM DTT.

GB1-His6-GluN2B(1170-1482) was purified with nickel-NTA agarose affinity column followed by size-exclusion chromatography using Superdex 200 (GE Healthcare) in a buffer containing 50 mM Tris pH 7.8, 300 mM NaCl, 1 mM EDTA, 1 mM DTT. The GB1-His6-tag was cleaved by HRV 3C protease and removed by another step of Superdex 200 size-exclusion chromatography using the same column buffer.

### Protein validation: LC-MS and UPLC

Protein concentrations were measured on a Nanodrop 1000 (Thermo Fisher Scientific) and all mass determinations were conducted on an electron spray ionization (ESI) liquid chromatography mass spectrometer (LC-MS) coupled to an Agilent 6410 triple quadrupole with two reverse phase C18 columns (Zorbax Eclipse XBD-C18, 4.6 × 50 mm) and (Agilent, Poroshell, 300SB-C18, 2.1 × 75 mm) for peptide and proteins respectively, using a binary buffer system consisting of H_2_O, Acetonitril (MeCN) and formic acid (buffer A: 95:5:0.1; buffer B: 5:95:0.1) at 0.75 mL/min. LC-MS samples were analyzed using the Agilent MassHunter version B.01.03 software. An analytical RP-UPLC (Waters Acquity) system with a reverse phase C18 column for peptides (Acquity UPLC BEH C18, 1.7 μm 2.1 × 50 mm) and C8 column for proteins (Acquity UPLC BEH C8, 1.7μM 2.1×50 mm) using a binary buffer system consisting of H2O, MeCN, trifluoroacetic acid (TFA) (buffer A: 95:5:0.1; buffer B: 5:95:0.1) was used to determine peptide and protein purity at 214 nm.

All samples for LC-MS and UPLC were diluted in buffer A (H_2_O, MeCN, TFA (95:5:0.1)) and a maximum of 2 μg protein was loaded.

### LC-MS/MS

#### Sample preparation

For the identification of site-specific serine phosphorylations, approximately 50 µg of protein in 50 mM NaPi pH 8 was subjected to sample preparation for LC-MS/MS analysis. Prior to reduction, PDZ1 pSer142 was denatured in 8M/2M Urea/Thiourea as the only sample. Hereafter, protein disulphides were reduced with 1 mM dithiothreitol (DTT) and free cysteine thiols alkylated with 5.5 mM chloroacetamide (CAA) for 60 min at room temperature (RT). Protein was pre-digested with 1 µg LysC per 100 µg protein for 3-4 hrs shaking at RT, LysC cleaving on the carboxyl side of lysine residues. Protein solutions were diluted in 50 mM ammonium bicarbonate (ABC) pH 8 and depending on the specific phosphorylation site, the protein solutions were either incubated ON at RT shaking with trypsin, cleaving on the carboxyl side of arginine and lysine residues, or LysC, using 1 µg enzyme per 100 µg protein of both enzymes. Hereafter, peptides were acidified to pH 2.5 with trifluoroacetic acid (TFA) and spun down for 50 min at 4700 rpm. Peptides were loaded onto StageTips, 4xC18 microcolumns (Empore Disk C18)(Rappsilber et al., 2003). The C18 StageTips were activated, washed and equilibrated in methanol, 80 % acetonitrile in 0.1 % formic acid (FA) and 0.1 % FA, respectively, before peptide loading. After peptide loading, the StageTips were washed in 0.1 % FA, spun dry and peptides directly eluted in 40 % acetonitrile in 0.1 % FA. Acetonitrile was evaporated by vacuum centrifugation and peptides dissolved in 13 µL 0.1 % FA and diluted 1:500 before mass spectrometric analysis.

#### Mass spectrometry analysis

Samples were analyzed using an EASY-nLC 1200 UHPLC system (Thermo Fisher Scientific) coupled to a Q Exactive HF-X mass spectrometer (Thermo Fisher Scientific). Separation of peptides was performed on a 15 cm analytical column with an inner diameter of 75 µm, packed in-house using ReproSil-Pur 120 C18-AQ 1.9 µm beads (Dr. Maisch). The analytical column was heated to 40 °C, and elution of peptides from the column was achieved by application of gradients with Buffer A (0.1 % formic acid) and increasing amounts of Buffer B (80 % acetonitrile in 0.1 % formic acid) at a flow rate of 250 nL/min. The primary gradient ranged from 5 % Buffer B to 45 % Buffer B over 30 min. followed by a wash block and equilibration. Electrospray ionization was achieved using a Nanospray Flex Ion Source (Thermo Fisher Scientific). Spray voltage was set to 2 kV, capillary temperature to 275 °C, and RF level to 40 %. Full scans were performed at a resolution of 60,000, a maximum injection time of 45 ms, with a scan range of 300 to 1,750 *m/z*, and an automatic gain control (AGC) target of 3e6 charges. Precursors were isolated with a width of 1.3 *m/z*, with an AGC target of 2e5 charges, and precursor fragmentation was accomplished using higher-energy collisional dissociation (HCD), with normalized collision energy of 28. Fragment scans were performed at a resolution of 60,000, a maximum injection time of 120 ms, a loop count of 7, and a scan range of 200 to 2,000 *m/z*. Selected precursors were excluded from repeated sequencing by setting a dynamic exclusion of 30 s.

#### Data analysis

All RAW files were analyzed using MaxQuant software (version 1.5.3.30). Default settings were used except as outlined below. For generation of the theoretical spectral library, a HUMAN.fasta database was extracted from UniProt on the 24^th^ of May, 2019. N-terminal acetylation, methionine oxidation, and phosphorylation (S, T, and Y) were set as variable modifications, and cysteine carbamidomethylation as a fixed modification. A maximum allowance of 5 variable modifications per peptide was used, and up to 5 missed cleavages was allowed. Mass tolerance for precursors was set to 20 ppm in the first MS/MS search and 4.5 ppm in the main MS/MS search after mass recalibration. For fragment masses, a tolerance of 20 ppm was used. Data was automatically filtered to achieve a false discovery rate of 1 % (default) at the peptide-spectrum match, the protein assignment, and the site-specific level. The output Phospho(S/T/Y)site table from MaxQuant was filtered to only include sites with a localization probability >0.99, a Score >100, and a Delta score >100.

### Circular dichroism

Circular dichroism (CD) experiments were made on a Jasco J-1500 spectrophotometer with a peltier controlled temperature. Prior to CD measurements, proteins were dialyzed ON at 4 °C into 50 mM NaPi, pH 7.4. The CD spectrums were measured on protein samples with concentrations of 15 µM, 10 µM or 1 µM for PSD-95 PDZ1, PSD-95 PDZ1-2 and PSD-95, respectively. Three accumulated scans were acquired for far-UV spectrum at 260-190 nm at 25 °C, and 95 °C directly followed by a 25 °C scan to assess the refolding of the protein, and at 25 °C in 6 M urea. Spectra for thermal denaturation were recorded at 260-190 nm from 20-95 °C with 5 °C intervals. All recorded CD data was obtained in millidegrees and converted to the molar ellipticity constant [θ], deg · cm^2^/dmol. The unit was converted based on the mean residual weight, pathlength and protein concentration. The data was analyzed using GraphPad Prism.

### Peptide synthesis

#### Manual peptide synthesis

Unless otherwise stated, amino acids and reagents were purchased from either Iris Biotech or Sigma Aldrich. Peptides were synthesized by SPPS using a 9-fluorenylmethyloxycarbonyl (Fmoc)-strategy at 0.1 mmol scale. Pre-loaded ChemMatrix resins (Sigma Aldrich) were used. Standard coupling reactions were achieved with 1:4:4:8 [resin: N-protected amino acid (AA): 2-(1-benzotriazole-1-yl)-1,1,3,3-tetramethyluronium hexafluorophosphate (HBTU), *N*,*N*-diisopropylethylamine (DIPEA)] or 1:4:4:8 [resin: AA: 1-(bis(dimethylamino)methylene]-1*H*-1,2,3-triazolo(4,5-b)pyridinium 3-oxid hexafluorophosphate (HATU): collidine]. Coupling reactions were agitated at room temperature (RT) and monitored using a Kaiser test kit (Sigma Aldrich). De-protections were carried out with 20 % piperidine (2 × 2 min). After coupling or de-protection steps the resin was extensively washed with dimethylformamide (DMF).

#### Automated peptide synthesis

Automated peptide synthesis using Fmoc-based SPPS was carried out on a Prelude X, induction heating assisted, peptide synthesizer (Gyros Protein Technologies, Tucson, AZ, USA) with 10 mL glass reaction vessel using preloaded Wang-resins (100–200 mesh). Reagents were prepared as solutions in DMF: Fmoc-protected AA (0.2 M), *O*-(1*H*-6-Chlorobenzotriazole-1-yl)-1,1,3,3-tetramethyluronium hexafluorophosphate (HCTU, 0.5 M), and DIPEA (1.0 M). Sequence elongation was achieved using the following protocol: deprotection (2 × 2 min, rt, 300 rpm shaking) and coupling (2 × 5 min, 75 °C, 300 rpm shaking, for Arg and His couplings 2 × 5 min, 50 °C, 300 rpm shaking). Amino acids were double coupled using amino acid/HCTU/DIPEA (ratio 1:1.25:2.5) in 5-fold excess over the resin loading to ensure efficient peptide elongation.

#### TAMRA coupling

N-terminal labelling of peptides with 5-(and-6)-carboxytetramethylrhodamine (TAMRA, Anaspec Inc.) was performed on resin, by coupling TAMRA for 16 h at rt using a mixture in NMP of TAMRA:benzotriazol-1-yloxy)tripyrrolidinophosphonium hexafluorophosphate (PyBOP):DIPEA (1.5:1.5:3) (Witte et al., 2013). To avoid photo bleaching of the fluorophore, the reaction vessel was covered and the coupling finalized with extensive resin washes with DMF and DCM.

#### Cleavage and purification

Peptides were cleaved from the resin using a mixture of 95:2.5:2.5 (trifluoroacetic acid (TFA), H_2_O and triisopropylsilane (TIPS)) for 2 hours at RT. After cleavage the peptides were precipitated in ice-cold diethyl ether. The precipitate was centrifuged 4350 rpm, 10 min, 4°C, washed with cold diethyl ether and the centrifugation step was repeated. The resulting peptide precipitate was dissolved in 50:50:0.1 (MeCN, H_2_O, TFA), filtered and lyophilized. All peptides were purified to >95% purity using a preparative RP-HPLC system with a reverse phase C18 column. A linear gradient from 0-40% using a binary buffer system of H_2_O, MeCN, TFA (buffer A: 95:5:0.1; buffer B: 5:95:0.1) at 20 mL/min was used. The purity of the collected fractions was measured at 214 nm on RP-UPLC at 0.45 mL/min and the final peptide products were lyophilized. The final products were further verified on LC-MS and the peptides diluted in 62.5 mM Tris-HCl pH 8 in and stored at −20°C.

### Fluorescence polarization

The fluorescence polarization (FP) assays were generally performed as previously described (Pedersen et al., 2017). In brief, binding affinities were measured using a Tecan Safire2 Microplate reader (Tecan) in flat bottom black 384-well plates (Corning Life Science) at 25 °C. The assays were conducted in 25 mM HEPES, 150 mM NaCl, pH 8, 1 mM TCEP. The TAMRA-labeled probes were measured at excitation/emission at 530/585 nm and the probe concentration was 200 nM in all FP experiments with C-terminal peptide ligands and 10 nM for AVLX-144. Prior to each measurement the instrumental Z-factor was adjusted to maximum fluorescence and the G-factor was calibrated to give an initial millipolarization (mP) at 20 in the probe reference well. For each binding setup, measurements were repeated 3 times. The obtained saturation curve was subtracted from the background and fitted to a one-sided binding model in GraphPad Prism.

### Phase transition sedimentation and imaging assay

Proteins were generally prepared as previously described (Zeng et al., 2018). Proteins were prepared in buffer containing 50mM Tris, pH 8.2, 100mM NaCl, 1 mM EDTA, and 2mM DTT and pre-cleared via high-speed centrifugations. Proteins were then mixed or diluted with buffer to designed combinations and concentrations. For sedimentation assay, typically, the final volume of each reaction is 100 µl. After 10 min equilibrium at room temperature, protein samples were subjected to sedimentation at 16,873 g for 10 min at 25⁰C on a table-top temperature-controlled micro-centrifuge.

After centrifugation, the supernatant and pellet were immediately separated into two tubes. The pellet fraction was thoroughly re-suspended with the same buffer to the equal volume as supernatant fraction (typically, to 100 µl). Proteins from both fractions were analyzed by SDS-PAGE with Coomassie blue staining. Band intensities were quantified using the ImageJ software. For imaging assay, protein samples were injected into a homemade flow chamber (comprised of a glass slide sandwiched by a coverslip with one layer of double-sided tape as a spacer) for DIC and fluorescent imaging (Nikon Ni-U upright fluorescence microscope) at room temperature. Glasses were washed by Hellmanex II (Hëlma Analytics) and 2 M NaOH sequentially and thoroughly rinsed with MilliQ H2O before chamber making. During imaging, the chamber was sealed by nail polish to reduce solution evaporation. Image fluorescence intensities were analyzed by the ImageJ software.

### Isothermal titration calorimetry (ITC) assay

ITC measurements were carried out on a Microcal VP-ITC calorimeter at 25 ⁰C. Proteins used for ITC measurements were dissolved in an assay buffer composed of 50 mM Tris, pH 8.2, 100 mM NaCl, 1 mM EDTA, and 2 mM DTT. High concentration of protein was loaded into the syringe and titrated into the cell containing low concentration of corresponding interactors (concentrations for each reaction were indicated in the figure legends). For each titration point, a 10 µL aliquot of a protein sample in the syringe was injected into the interacting protein in the cell at a time interval of 2 min. Titration data were analyzed using the Origin7.0 software and fitted with the one-site binding model.

## REFERENCES

Bach, A., Clausen, B. H., Moller, M., Vestergaard, B., Chi, C. N., Round, A., Sorensen, P. L., Nissen, K. B., Kastrup, J. S., Gajhede, M., Jemth, P., Kristensen, A. S., Lundstrom, P., Lambertsen, K. L. & Stromgaard, K. 2012. A High-affinity, dimeric inhibitor of PSD-95 bivalently interacts with PDZ1-2 and protects against ischemic brain damage. Proc Natl Acad Sci U S A, 109, 3317–22.

Ballif, B. A., Carey, G. R., Sunyaev, S. R. & Gygi, S. P. 2008. Large-Scale Identification and Evolution Indexing of Tyrosine Phosphorylation Sites from Murine Brain. Journal of Proteome Research, 7, 311–318.

Banani, S. F., Lee, H. O., Hyman, A. A. & Rosen, M. K. 2017. Biomolecular condensates: organizers of cellular biochemistry. Nat Rev Mol Cell Biol, 18, 285–298.

Barber, K. W., Muir, P., Szeligowski, R. V., Rogulina, S., Gerstein, M., Sampson, J. R., Isaacs, F. J. & Rinehart, J. 2018. Encoding human serine phosphopeptides in bacteria for proteome-wide identification of phosphorylation-dependent interactions. Nat Biotechnol, 36, 638–644.

Chen, L., Chetkovich, D. M., Petralia, R. S., Sweeney, N. T., Kawasaki, Y., Wenthold, R. J., Bredt, D. S. & Nicoll, R. A. 2000. Stargazin regulates synaptic targeting of AMPA receptors by two distinct mechanisms. Nature, 408, 936–943.

Chen, X., Levy, J. M., Hou, A., Winters, C., Azzam, R., Sousa, A. A., Leapman, R. D., Nicoll, R. A. & Reese, T. S. 2015. PSD-95 family MAGUKs are essential for anchoring AMPA and NMDA receptor complexes at the postsynaptic density. Proc Natl Acad Sci U S A, 112, E6983–92.

Chen, X., Nelson, C. D., Li, X., Winters, C. A., Azzam, R., Sousa, A. A., Leapman, R. D., Gainer, H., Sheng, M. & Reese, T. S. 2011. PSD-95 is required to sustain the molecular organization of the postsynaptic density. J Neurosci, 31, 6329–38.

Chen, X., Wu, X., Wu, H. & Zhang, M. 2020. Phase separation at the synapse. Nat Neurosci.

Chetkovich, D. M., Chen, L., Stocker, T. J., Nicoll, R. A. & Bredt, D. S. 2002. Phosphorylation of the Postsynaptic Density-95 (PSD-95)/Discs Large/Zona Occludens-1 Binding Site of Stargazin Regulates Binding to PSD-95 and Synaptic Targeting of AMPA Receptors. The Journal of Neuroscience, 22, 5791–5796.

Chi, C. N., Bach, A., Stromgaard, K., Gianni, S. & Jemth, P. 2012. Ligand binding by PDZ domains. Biofactors, 38, 338–48.

Chin, J. W. 2017. Expanding and reprogramming the genetic code. Nature, 550, 53–60.

Coley, A. A. & Gao, W. J. 2018. PSD95: A synaptic protein implicated in schizophrenia or autism? Prog Neuropsychopharmacol Biol Psychiatry, 82, 187–194.

Doyle, D. A., Lee, A., Lewis, J., Kim, E., Sheng, M. & Mackinnon, R. 1996. Crystal Structures of a Complexed and Peptide-Free Membrane Protein-Binding Domain: Molecular Basis of Peptide Recognition by PDZ. Cell, 85, 1067–1076.

Du, C. P., Gao, J., Tai, J. M., Liu, Y., Qi, J., Wang, W. & Hou, X. Y. 2009. Increased tyrosine phosphorylation of PSD-95 by Src family kinases after brain ischaemia. Biochem J, 417, 277–85.

El-Husseini, A. E.-D., Schnell, E., Chetkovich, D. M., Nicoll, R. A. & Bredt, D. S. 2000. PSD-95 Involvement in Maturation of Excitatory Synapses. Science, 290, 1364–1368.

Feng, W. & Zhang, M. 2009. Organization and dynamics of PDZ-domain-related supramodules in the postsynaptic density. Nat Rev Neurosci, 10, 87–99.

Funke, L., Dakoji, S. & Bredt, D. S. 2005. Membrane-associated guanylate kinases regulate adhesion and plasticity at cell junctions. Annu Rev Biochem, 74, 219–45.

Gardoni, F., Polli, F., Cattabeni, F. & Di Luca, M. 2006. Calcium-calmodulin-dependent protein kinase II phosphorylation modulates PSD-95 binding to NMDA receptors. Eur J Neurosci, 24, 2694–704.

Gray, E. G. 1959. Axo-somatic and axo-dendritic synapses of the cerebral cortex: an electron microscope study. Jourrnal of Anatomy, 93, 420–433.

Harris, B. Z., Hillier, B. J. & Lim, W. A. 2001. Energetic Determinants of Internal Motif Recognition by PDZ Domains. Biochemistry, 40, 5921–5930.

Heinemann, I. U., Rovner, A. J., Aerni, H. R., Rogulina, S., Cheng, L., Olds, W., Fischer, J. T., Soll, D., Isaacs, F. J. & Rinehart, J. 2012. Enhanced phosphoserine insertion during Escherichia coli protein synthesis via partial UAG codon reassignment and release factor 1 deletion. FEBS Lett, 586, 3716–22.

Hill, M. D., Martin, R. H., Mikulis, D., Wong, J. H., Silver, F. L., Terbrugge, K. G., Milot, G., Clark, W. M., Macdonald, R. L., Kelly, M. E., Boulton, M., Fleetwood, I., Mcdougall, C., Gunnarsson, T., Chow, M., Lum, C., Dodd, R., Poublanc, J., Krings, T., Demchuk, A. M., Goyal, M., Anderson, R., Bishop, J., Garman, D. & Tymianski, M. 2012. Safety and efficacy of NA-1 in patients with iatrogenic stroke after endovascular aneurysm repair (ENACT): a phase 2, randomised, double-blind, placebo-controlled trial. The Lancet Neurology, 11, 942–950.

Hyman, A. A., Weber, C. A. & Julicher, F. 2014. Liquid-liquid phase separation in biology. Annu Rev Cell Dev Biol, 30, 39–58.

Kim, E., Niethammer, M., Rothschild, A., Jan, Y. N. & Sheng, M. 1995. Clustering of Shaker-type K+ channels by interaction with a family of membraneÂ· associated guanylate kinases. Nature, 378, 85–88.

Kim, E. & Sheng, M. 2004. PDZ domain proteins of synapses. Nat Rev Neurosci, 5, 771–81.

Kornau, H.-C., Schenker, L. T., Kennedy, M. B. & Seeburg, P. H. 1995. Domain Interaction between NMDA Receptor Subunits and the Postsynaptic DensityProtein PSD-95. Science, 269, 1737–1740.

Lajoie, M. J., Rovner, A. J., Goodman, D. B., Aerni, H. R., Haimovich, A. D., Kuznetsov, G., Mercer, J. A., Wang, H. H., Carr, P. A., Mosberg, J. A., Rohland, N., Schultz, P. G., Jacobson, J. M., Rinehart, J., Church, G. M. & Isaacs, F. J. 2013. Genomically recoded organisms expand biological functions. Science, 342, 357–60.

Lee, H. J. & Zheng, J. J. 2010. PDZ domains and their binding partners: structure, specificity, and modification. Cell Commun Signal, 8, 8.

Lee, S., Oh, S., Yang, A., Kim, J., Soll, D., Lee, D. & Park, H. S. 2013. A facile strategy for selective incorporation of phosphoserine into histones. Angew Chem Int Ed Engl, 52, 5771–5.

Li, P., Banjade, S., Cheng, H. C., Kim, S., Chen, B., Guo, L., Llaguno, M., Hollingsworth, J. V., King, D. S., Banani, S. F., Russo, P. S., Jiang, Q. X., Nixon, B. T. & Rosen, M. K. 2012. Phase transitions in the assembly of multivalent signalling proteins. Nature, 483, 336–40.

Li, Q., Peng, X., Li, Y., Tang, W., Zhu, J., Huang, J., Qi, Y. & Zhang, Z. 2020. LLPSDB: a database of proteins undergoing liquid-liquid phase separation in vitro. Nucleic Acids Res, 48, D320–D327.

Morabito, M. A., Sheng, M. & Tsai, L. H. 2004. Cyclin-dependent kinase 5 phosphorylates the N-terminal domain of the postsynaptic density protein PSD-95 in neurons. J Neurosci, 24, 865–76.

Mukai, T., Hoshi, H., Ohtake, K., Takahashi, M., Yamaguchi, A., Hayashi, A., Yokoyama, S. & Sakamoto, K. 2015. Highly reproductive Escherichia coli cells with no specific assignment to the UAG codon. Sci Rep, 5, 9699.

Nicoll, R. A., Tomita, S. & Bredt, D. S. 2006. Auxiliary Subunits Assist AMPA-Type Glutamate Receptors. Science, 311, 1253.

Niethammer, M., Kim, E. & Sheng, M. 1996. Interaction between the C terminus of NMDA receptor subunits and multiple members of the PSD-95 family of membrane-associated guanylate kinases. The Journal of Neuroscience, 16, 2157–2163.

Nishiyama, J. & Yasuda, R. 2015. Biochemical Computation for Spine Structural Plasticity. Neuron, 87, 63–75.

Park, H. S., Hohn, M. J., Umehara, T., Guo, L. T., Osborne, E. M., Benner, J., Noren, C. J., Söll, D. & Rinehart, J. 2011. Expanding the Genetic Code of Escherichia coli with Phosphoserine. Science, 33.

Pedersen, S. W., Albertsen, L., Moran, G. E., Levesque, B., Pedersen, S. B., Bartels, L., Wapenaar, H., Ye, F., Zhang, M., Bowen, M. E. & Stromgaard, K. 2017. Site-Specific Phosphorylation of PSD-95 PDZ Domains Reveals Fine-Tuned Regulation of Protein-Protein Interactions. ACS Chem Biol, 12, 2313–2323.

Pedersen, S. W., Pedersen, S. B., Anker, L., Hultqvist, G., Kristensen, A. S., Jemth, P. & Stromgaard, K. 2014. Probing backbone hydrogen bonding in PDZ/ligand interactions by protein amide-to-ester mutations. Nat Commun, 5, 3215.

Pirman, N. L., Barber, K. W., Aerni, H. R., Ma, N. J., Haimovich, A. D., Rogulina, S., Isaacs, F. J. & Rinehart, J. 2015. A flexible codon in genomically recoded Escherichia coli permits programmable protein phosphorylation. Nat Commun, 6, 8130.

Rao, A., Kim, E., Sheng, M. & Craig, A. M. 1998. Heterogeneity in the Molecular Composition of Excitatory Postsynaptic Sites during Development of Hippocampal Neurons in Culture. The Journal of Neuroscience, 18, 1217–1229.

Rappsilber, J., Ishihama, Y. & Mann, M. 2003. Stop and Go Extraction Tips for Matrix-Assisted Laser Desorption/Ionization, Nanoelectrospray, and LC/MS Sample Pretreatment in Proteomics. Analytical Chemistry, 75, 663–670.

Rogerson, D. T., Sachdeva, A., Wang, K., Haq, T., Kazlauskaite, A., Hancock, S. M., Huguenin-Dezot, N., Muqit, M. M., Fry, A. M., Bayliss, R. & Chin, J. W. 2015. Efficient genetic encoding of phosphoserine and its nonhydrolyzable analog. Nat Chem Biol, 11, 496–503.

Sattler, R., Xiong, Z., Lu, W. Y., Hafner, M., Macdonald, J. F. & Tymiansky, M. 1999. Specific Coupling of NMDA Receptor Activation to Nitric Oxide Neurotoxicity by PSD-95 Protein. Science, 284, 1845–1848.

Schwark, D. G., Schmitt, M. A. & Fisk, J. D. 2018. Dissecting the Contribution of Release Factor Interactions to Amber Stop Codon Reassignment Efficiencies of the Methanocaldococcus jannaschii Orthogonal Pair. Genes (Basel), 9.

Sheng, M. & Hoogenraad, C. C. 2007. The postsynaptic architecture of excitatory synapses: a more quantitative view. Annu Rev Biochem, 76, 823–47.

Sheng, M. & Kim, E. 2011. The postsynaptic organization of synapses. Cold Spring Harb Perspect Biol, 3.

Vallejo, D., Codocedo, J. F. & Inestrosa, N. C. 2016. Posttranslational Modifications Regulate the Postsynaptic Localization of PSD-95. Mol Neurobiol.

Volk, L., Chiu, S. L., Sharma, K. & Huganir, R. L. 2015. Glutamate synapses in human cognitive disorders. Annu Rev Neurosci, 38, 127–49.

Witte, M. D., Theile, C. S., Wu, T., Guimaraes, C. P., Blom, A. E. & Ploegh, H. L. 2013. Production of unnaturally linked chimeric proteins using a combination of sortase-catalyzed transpeptidation and click chemistry. Nat Protoc, 8, 1808–19.

Xie, J. & Schultz, P. G. 2005. An expanding genetic code. Methods, 36, 227–38.

Xu, H., Wang, Y., Lu, J., Zhang, B., Zhang, Z., Si, L., Wu, L., Yao, T., Zhang, C., Xiao, S., Zhang, L., Xia, Q. & Zhou, D. 2016. Re-exploration of the Codon Context Effect on Amber Codon-Guided Incorporation of Noncanonical Amino Acids inEscherichia coliby the Blue-White Screening Assay. ChemBioChem, 17, 1250–1256.

Xue, Y., Ren, J., Gao, X., Jin, C., Wen, L. & Yao, X. 2008. GPS 2.0, a tool to predict kinase-specific phosphorylation sites in hierarchy. Mol Cell Proteomics, 7, 1598–608.

Ye, F. & Zhang, M. 2013. Structures and target recognition modes of PDZ domains: recurring themes and emerging pictures. Biochem J, 455, 1–14.

Zeng, M., Chen, X., Guan, D., Xu, J., Wu, H., Tong, P. & Zhang, M. 2018. Reconstituted Postsynaptic Density as a Molecular Platform for Understanding Synapse Formation and Plasticity. Cell, 174, 1172–1187 e16.

Zeng, M., Diaz-Alonso, J., Ye, F., Chen, X., Xu, J., Ji, Z., Nicoll, R. A. & Zhang, M. 2019. Phase Separation-Mediated TARP/MAGUK Complex Condensation and AMPA Receptor Synaptic Transmission. Neuron, 104, 529–543 e6.

Zeng, M., Shang, Y., Araki, Y., Guo, T., Huganir, R. L. & Zhang, M. 2016. Phase Transition in Postsynaptic Densities Underlies Formation of Synaptic Complexes and Synaptic Plasticity. Cell, 166, 1163–1175 e12.

Zhu, J., Shang, Y. & Zhang, M. 2016. Mechanistic basis of MAGUK-organized complexes in synaptic development and signalling. Nat Rev Neurosci, 17, 209–23.

Zhu, P., Gafken, P. R., Mehl, R. A. & Cooley, R. B. 2019. A Highly Versatile Expression System for the Production of Multiply Phosphorylated Proteins. ACS Chem Biol, 14, 1564–1572.

Aarts, M., Liu, Y., Liu, L., Besshoh, S., Arundine, M., Gurd, J. W., Wang, Y. T., Salter, M. W. & Tymianski, M. 2002. Treatment of Ischemic Brain Damage by Perturbing NMDA Receptor-PSD-95 Protein Interactions. Science, 298, 846–850.

