## Supporting Information for "Site-specific phosphorylation of PSD-95 dynamically regulates the post-synaptic density as observed by phase separation"

### Supplemental Information

#### SUPPLEMENTARY FIGURES WITH TITLES AND LEGENDS

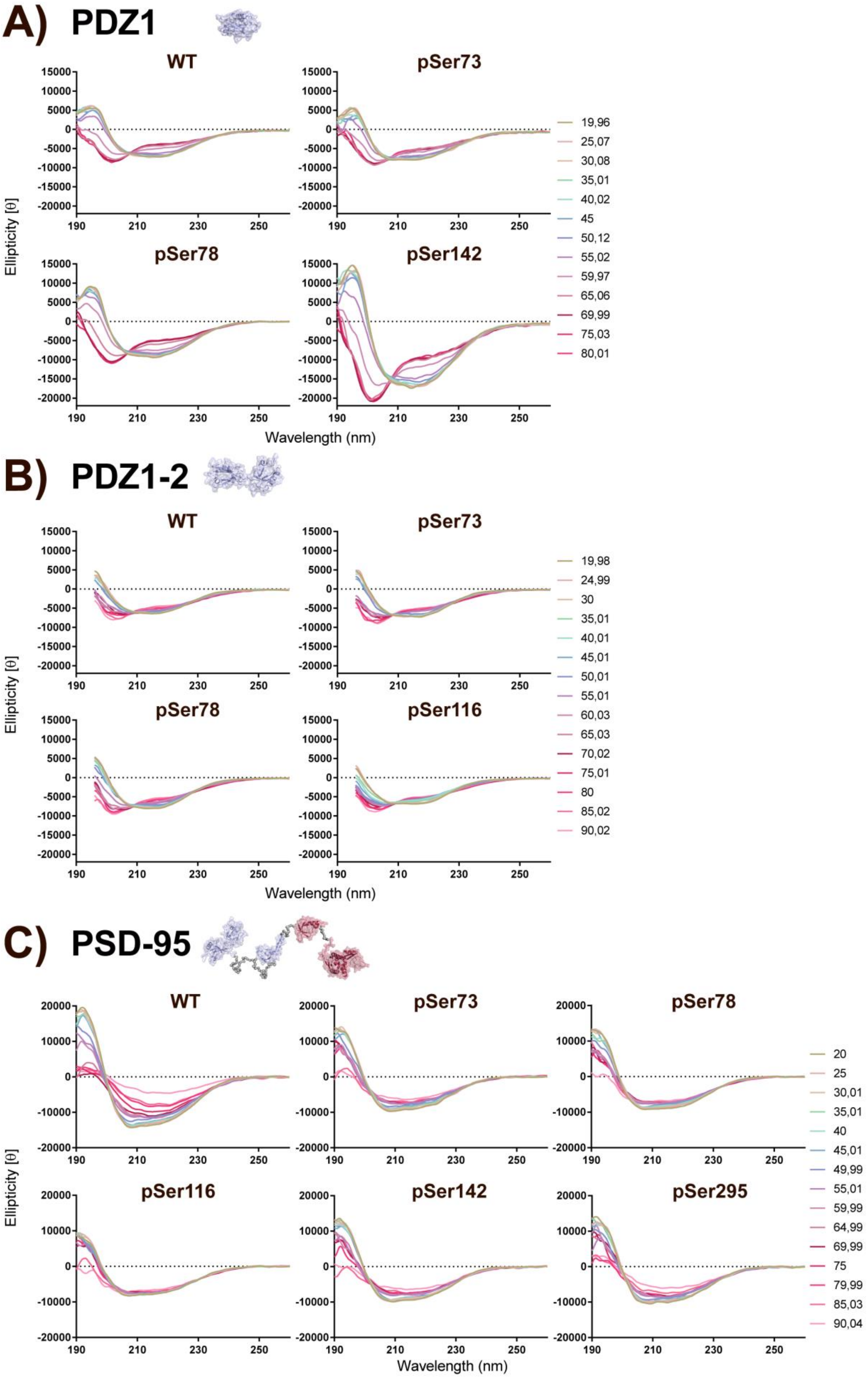
A) PDZ1
WT
pSer73
pSer78
pSer142
Ellipticity [θ]
Wavelength (nm)
19,96
25,07
30,08
35,01
40,02
45
50,12
55,02
59,97
65,06
69,99
75,03
80,01
B) PDZ1-2
WT
pSer73
pSer78
pSer116
Ellipticity [θ]
Wavelength (nm)
19,98
24,99
30
35,01
40,01
45,01
50,01
55,01
60,03
65,03
70,02
75,01
80
85,02
90,02
C) PSD-95
WT
pSer73
pSer78
pSer116
pSer142
pSer295
Ellipticity [θ]
Wavelength (nm)
20
25
30,01
35,01
40
45,01
49,99
55,01
59,99
64,99
69,99
75
79,99
85,03
90,04

**Supplementary Figure 1: Thermal denaturation CD spectra of all protein variants**

Thermal denaturation CD spectra of all protein variants of A) PDZ1, B) PDZ1-2 and C) PSD-95. Scans were obtained at 260-190 nm from 20-90 $^{o}$C with 5 $^{o}$C intervals, except for PDZ1, where unfolding was measured from 20-80 $^{o}$C. Data shown does not exceed a high tension voltage above 700 V.

### PDZ1

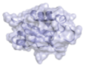

## A) WT

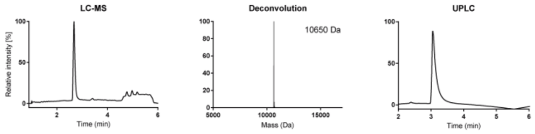


#### B) pSer73

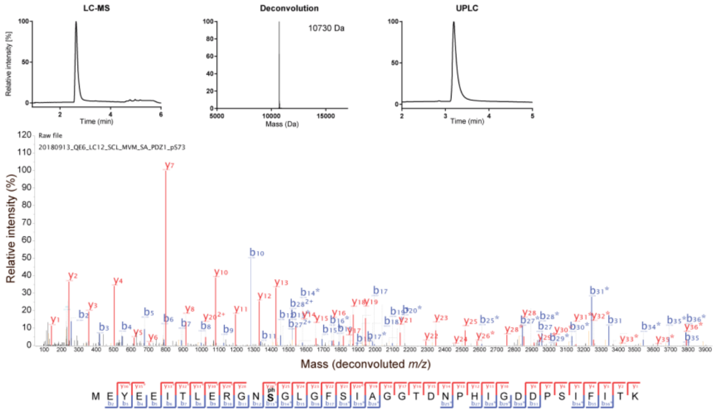


#### C) pSer78

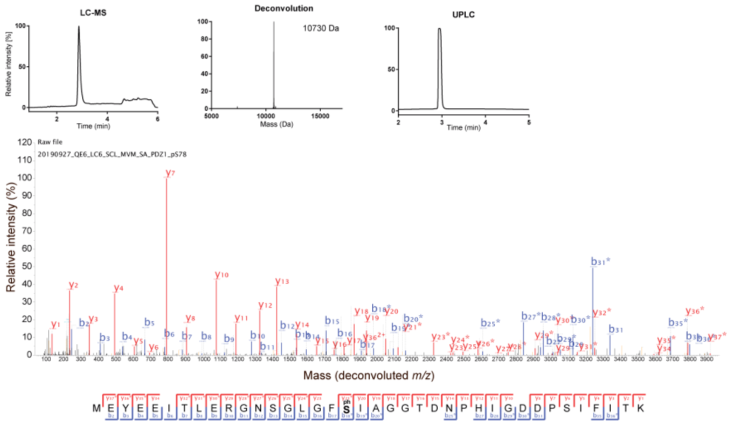


#### D) pSer142

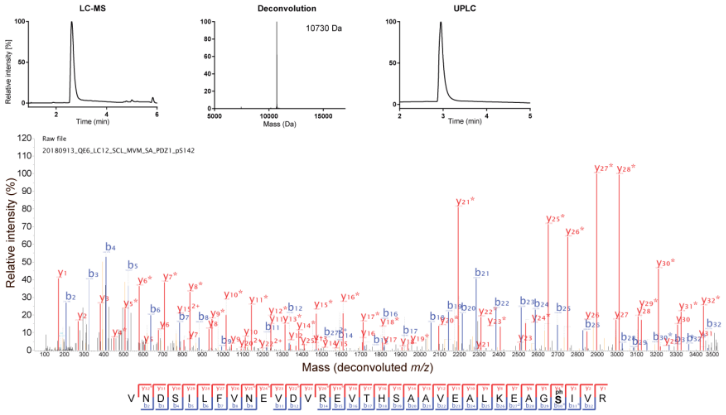

**Supplementary Figure 2: LCMS, UPLC and LC-MS/MS results for PDZ1**

For purified the protein variants of PDZ1, LC-MS spectra and masses derived from the deconvoluted spectra, and UPLC chromatograms are depicted on top, and the annotated mass spectrums and identified peptides sequences below. The identified phosphorylated Ser residue is highlighted. Full fragmentation is obtained with both b-ions (blue) and y-ions (red) indicated. **A)** WT, **B)** pSer73, **C)** pSer78 and **D)** pSer142. The expected masses for PDZ1 were 10650 Da and 10730 Da for the WT and pSer protein variants, respectively.

.

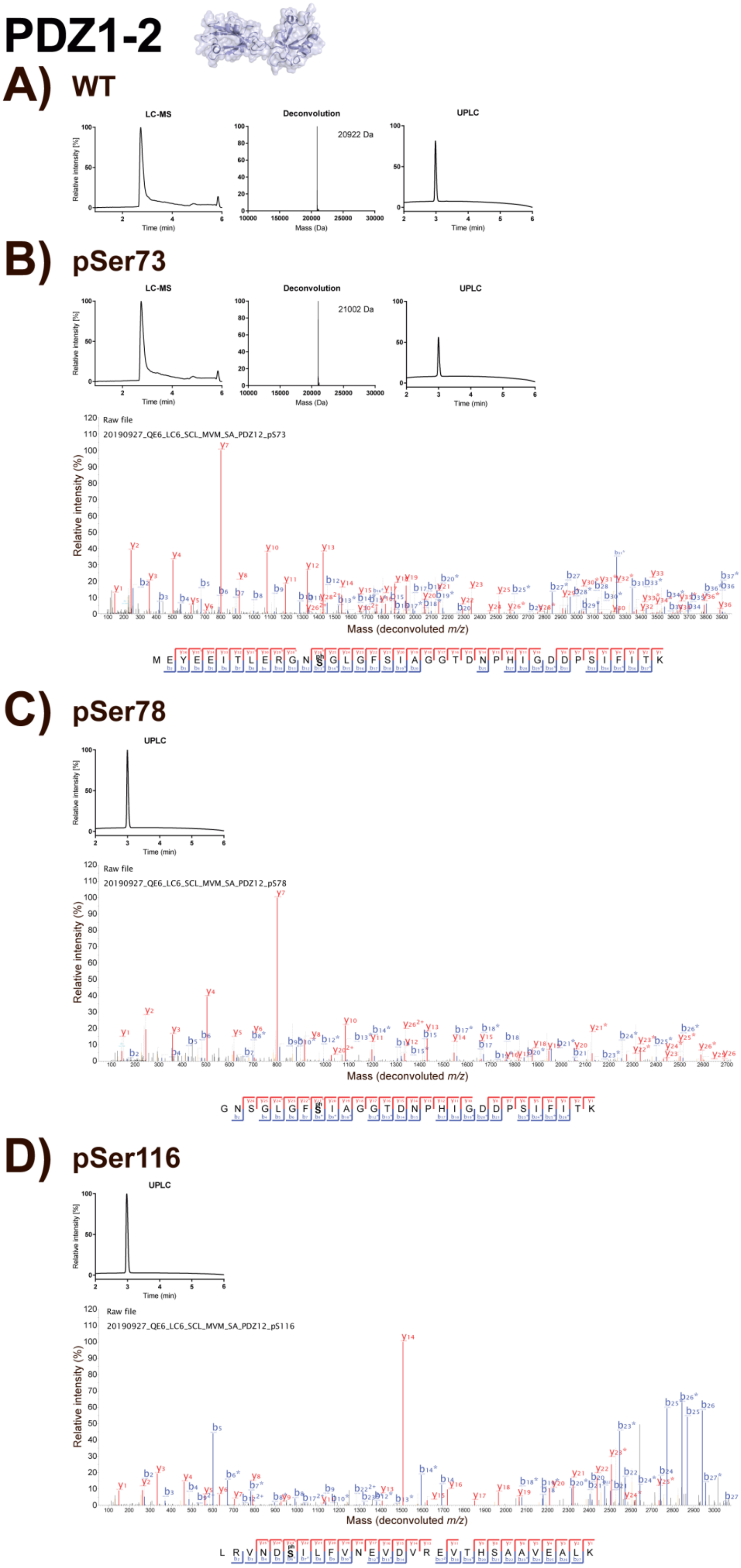

PDZ1-2
A) WT
LC-MS
Deconvolution
UPLC
20922 Da
Relative intensity [%]
Time (min)
Mass (Da)
B) pSer73
LC-MS
Deconvolution
UPLC
21002 Da
Raw file
20190927_QE6_LC6_SCL_MVM_SA_PDZ12_pS73
Relative intensity (%)
Mass (deconvoluted m/z)
M E Y E E I T L E R G N S G L G F S I A G G T D N P H I G D D P S I F I T K
C) pSer78
UPLC
Raw file
20190927_QE6_LC6_SCL_MVM_SA_PDZ12_pS78
Relative intensity (%)
Mass (deconvoluted m/z)
G N S G L G F S I A G G T D N P H I G D D P S I F I T K
D) pSer116
UPLC
Raw file
20190927_QE6_LC6_SCL_MVM_SA_PDZ12_pS116
Relative intensity (%)
Mass (deconvoluted m/z)
L R V N D S I L F V N E V D V R E V T H S A A V E A L K

**Supplementary Figure 3: LCMS, UPLC and LC-MS/MS for PDZ1-2**

For purified the protein variants of PDZ1-2, LC-MS spectra and masses derived from the deconvoluted spectra, and UPLC chromatograms are depicted on top, and the annotated mass spectrums and identified peptides sequences below. The identified phosphorylated Ser residue is highlighted. Full fragmentation is obtained with both b-ions (blue) and y-ions (red) indicated. **A)** WT, **B)** pSer73, **C)** pSer78 and **D)** pSer116. The expected masses for PDZ1-2 were 20924 Da and 21004 Da for the WT and pSer protein variants, respectively.

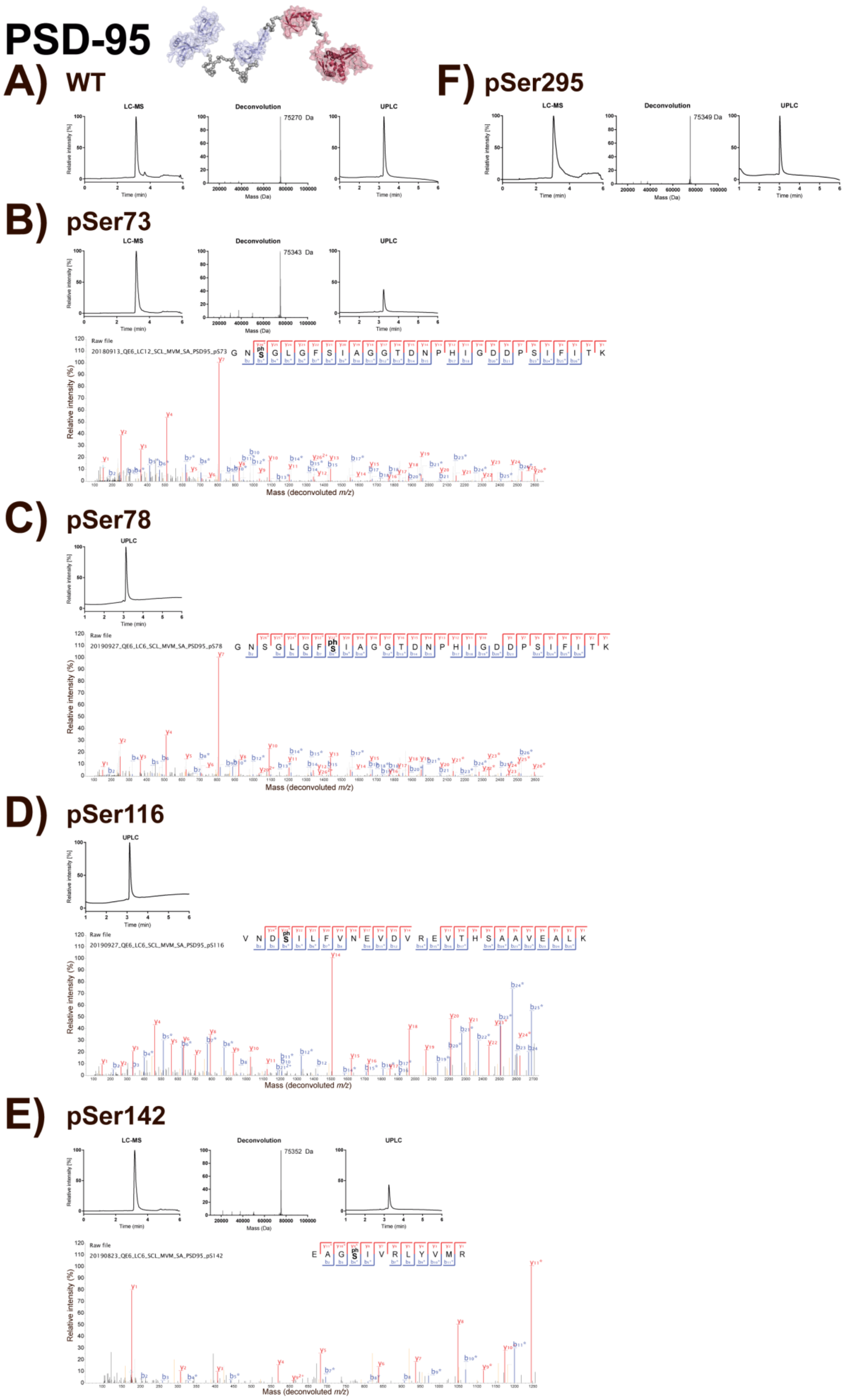

PSD-95
A) WT
LC-MS
Deconvolution
75270 Da
UPLC
F) pSer295
75349 Da
B) pSer73
75343 Da
Raw file
20180913_QE6_LC12_SCL_MVM_SA_PSD95_pS73
G N S G L G F S I A G G T D N P H I G D D P S I F I T K
Relative intensity (%)
Mass (deconvoluted m/z)
C) pSer78
20190927_QE6_LC6_SCL_MVM_SA_PSD95_pS78
D) pSer116
20190927_QE6_LC6_SCL_MVM_SA_PSD95_pS116
V N D S I L F V N E V D V R E V T H S A A V E A L K
E) pSer142
75352 Da
20190823_QE6_LC6_SCL_MVM_SA_PSD95_pS142
E A G S I V R L Y V M R
Relative intensity [%]
Time (min)
Mass (Da)

**Supplementary Figure 4: LCMS, UPLC and LC-MS/MS results for PSD-95**

For purified the protein variants of PSD-95, LC-MS spectra and masses derived from the deconvoluted spectra, and UPLC chromatograms are depicted on top, and the annotated mass spectrums and identified peptides sequences below. The identified phosphorylated Ser residue is highlighted. Full fragmentation is obtained with both b-ions (blue) and y-ions (red) indicated. **A)** WT, **B)** pSer73, **C)** pSer78, **D)** pSer116, **E)** pSer142 and **F)** pSer295. The expected masses for PDZ1-2 were 20924 Da and 21004 Da for the WT and pSer protein variants, respectively. The expected masses for PSD-95 were 75269 Da and 75349 Da for the WT and pSer protein variants, respectively.

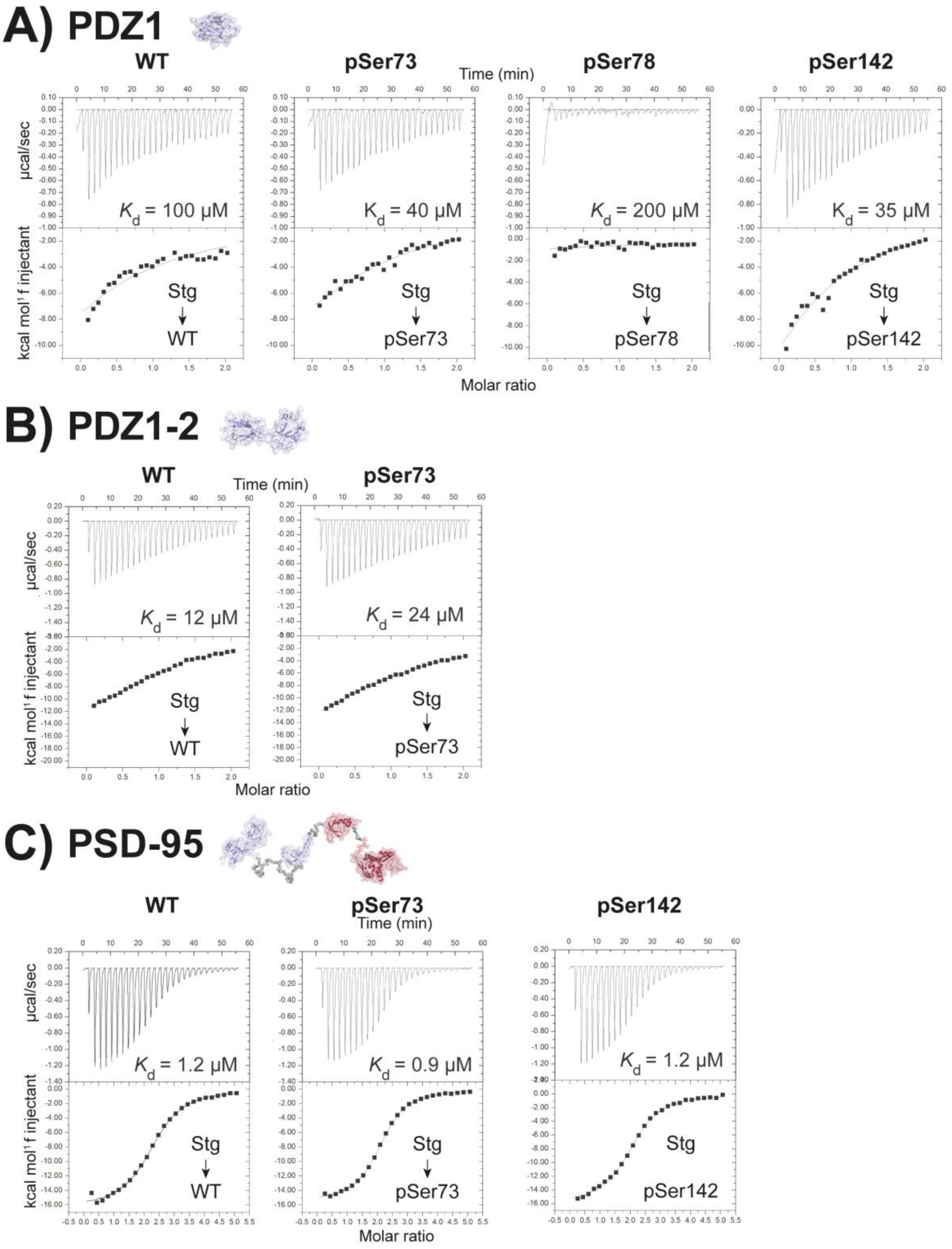

A) PDZ1
WT
pSer73
pSer78
pSer142
Time (min)
µcal/sec
kcal mol⁻¹ f injectant
$K_d$ = 100 µM
$K_d$ = 40 µM
$K_d$ = 200 µM
$K_d$ = 35 µM
Stg → WT
Stg → pSer73
Stg → pSer78
Stg → pSer142
Molar ratio
B) PDZ1-2
WT
pSer73
Time (min)
$K_d$ = 12 µM
$K_d$ = 24 µM
Stg → WT
Stg → pSer73
Molar ratio
C) PSD-95
WT
pSer73
pSer142
Time (min)
$K_d$ = 1.2 µM
$K_d$ = 0.9 µM
$K_d$ = 1.2 µM
Stg → WT
Stg → pSer73
Stg
pSer142
Molar ratio

**Supplementary Figure 5: ITC assays of protein variants**

**A)** ITC-based measurements of the PBM and C-terminal of stargazin (Stg) binding PDZ1 WT, pSer73, pSer78 and pSer142. 300 µM Stg was titrated into 30 µM protein in the cell and binding constants ($K_d$) were calculated.

**B)** ITC-based measurements of the PBM and C-terminal of stargazin (Stg) binding PDZ1-2 pSer73. 250 µM Stg was titrated into 25 µM protein in the cell and binding constants (*K*d) were calculated.

**C)** ITC-based measurements of the PBM and C-terminal of stargazin (Stg) binding PSD-95 pSer73 and pSer142. 250 µM Stg was titrated into 10 µM protein in the cell and binding constants ($K_d$) were calculated.

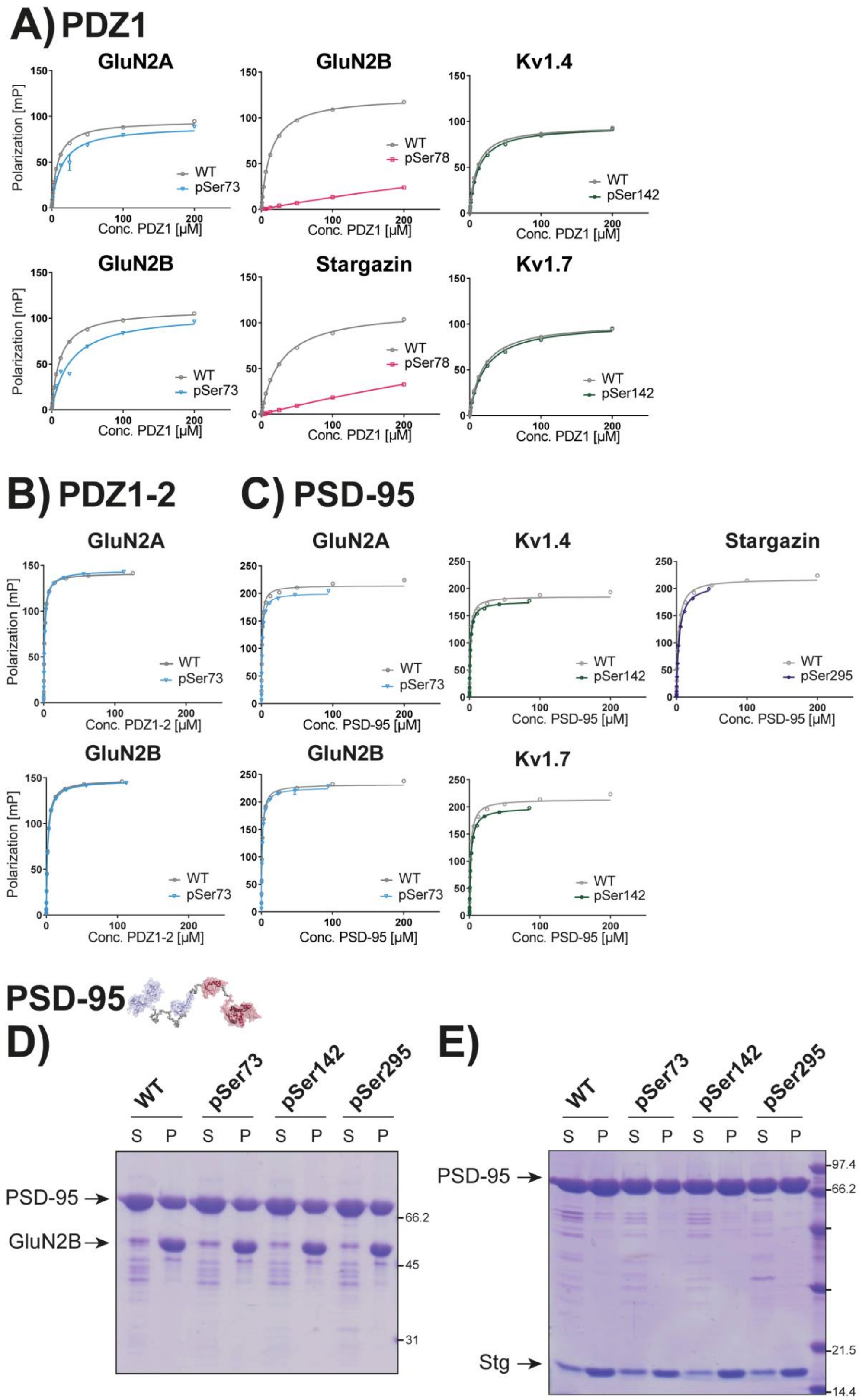

A) PDZ1
GluN2A
GluN2B
Kv1.4
Stargazin
Kv1.7
Polarization [mP]
Conc. PDZ1 [µM]
WT
pSer73
pSer78
pSer142
B) PDZ1-2
C) PSD-95
Conc. PDZ1-2 [µM]
Conc. PSD-95 [µM]
pSer295
PSD-95
D)
E)
WT
pSer73
pSer142
pSer295
S P S P S P S P
PSD-95 →
GluN2B →
Stg →
97.4
66.2
45
31
21.5
14.4

**Supplementary Figure 6: FP and sedimentation assays of protein variants**

**A)** PDZ1 protein variants were tested for binding to peptide ligands in a FP saturation assay. Here, 200 nM of TAMRA-labeled C-terminal peptide ligands were saturated with protein in a concentration dependent manner with a maximum concentration of 200 µM protein. The milli-polarization (mP) was measured for all concentrations in triplicates, background fluorescence subtracted and the data fitted to a one-sided binding model from which $K_d$ values were calculated (see **Supplementary table 4**). First row (top + bottom), PDZ1 pSer73 was tested for binding to GluN2A (top) and GluN2B (bottom). Second row (top + bottom), PDZ1 pSer78 was tested for binding to GluN2B (top) and stargazin (bottom). Third row (top + bottom), PDZ1 pSer142 was tested for binding to Kv1.4 (top) and Kv1.7 (bottom).

**B)** PDZ1-2 pSer73 protein variant was tested for binding to GluN2A (top) and GluN2B (bottom) peptide ligands in a FP saturation assay using the same conditions as in **A)**.

**C)** PSD-95 protein variants were tested for binding to peptide ligands in a FP saturation assay using the same conditions as in **A)**. First row (top + bottom), PSD-95 pSer73 was tested for binding to GluN2A (top) and GluN2B (bottom). Second row (top + bottom), PSD-95 pSer142 was tested for binding to Kv1.4 (top) and Kv1.7 (bottom). Third row (top + bottom), PSD-95 pSer295 was tested for binding to stargazin.

**D)** Representative SDS-PAGE data of sedimentation experiments showing the distributions of various forms of phosphorylated PSD-95 and GluN2B_CT recovered from the aqueous phase or supernatant (S) and the condensed phase or pellet (P). 10 µM GluN2B_CT and 5 µM FL PSD-95 was used for GluN2B-PSD-95 LLPS.

**E)** Representative SDS-PAGE data of sedimentation experiments showing the distributions of various forms of phosphorylated PSD-95 and Stg_CT recovered from the aqueous phase or supernatant (S) and the condensed phase or pellet (P). 10 μM Stg_CT and 10 μM PSD-95 was used for Stg-PSD-95 LLPS.

#### SUPPLEMENTARY TABLES WITH TITLES AND LEGENDS

#### Supplementary Table 1: Primers

| Primer name | Sequence | Application |
|---|---|---|
| MVM_034 | CGGCGGCCATGGAATACGAGGAAATCACATTGG | |
| MVM_035 | CCGCCGCCTAGGTCAATGATGATGATGATGGTGCCGGCGCATGACATAGA | |
| MVM_001 | GGAAAGGGGTAACTAGGGTCTGGGCTTCAGC | Ser73TAG forward |
| MVM_002 | GCTGAAGCCCAGACCCTAGTTACCCCTTTCC | Ser73TAG reverse |
| TLJ_001 | GGTCTGGGCTTCTAGATCGCAGGTGGC | Ser78TAG forward |
| TLJ_002 | GCCACCTGCGATCTAGAAGCCCAGACC | Ser78TAG reverse |
| TLJ_003 | CGGTGACGACCCATAGATTTTCATCACC | Ser93TAG forward |
| TLJ_004 | GGTGATGAAAATCTATGGGTCGTCACCG | Ser93TAG reverse |
| TLJ_005 | GGGTCAACGACTAGATCCTGTTTGT | Ser116TAG forward |
| TLJ_006 | ACAAACAGGATCTAGTCGTTGACCC | Ser116TAG reverse |
| TLJ_007 | GGTGACCCACTAGGCGGCGGTGGAA | Ser131TAG forward |
| TLJ_008 | TTCCACCGCCGCCTAGTGGGTCACC | Ser131TAG reverse |
| MVM_003 | CCTCAAAGAGGCAGGCTAGATCGTTCGCC | Ser142TAG forward |
| MVM_004 | GGCGAACGATCTAGCCTGCCTCTTTGAGG | Ser142TAG reverse |
| MVM_005 | CCCTCGGCGCTACTAGCCAGTGGCC | Ser295TAG forward |
| MVM_006 | GGCCACTGGCTAGTAGCGCCGAGGG | Ser295TAG reverse |
| MVM_036 | GCGTCCCTGCGGTAGAACCC | Ser425TAG forward |
| MVM_037 | GGGTTCTACCGCAGGGACGC | Ser425TAG reverse |
| TLJ_013 | CCCGACAAGTTTGGATAGTGTGTTCCCC | Ser561TAG forward |
| TLJ_014 | GGGGAACACACTATCCAAACTTGTCGGG | Ser561TAG reverse |

**Supplementary Table 2: $K_d$ values from FP assays with AVLX-144 (values in nM)**

| Protein | AVLX-144 |
|---|---|
| **PDZ1-2** | |
| **WT** | 16.76 ± 0.5 |
| **pSer73** | 37.63 ± 0.9 |
| **pSer78** | N.B. |
| **pSer116** | 41.68 ± 1.5 |
| **PSD-95** | |
| **WT** | 116.9 ± 9.8 |
| **pSer73** | 185.8 ± 31.82 |
| **pSer78** | N.B. |
| **pSer116** | 245 ± 44.6 |
| **pSer142** | 208.8 ± 19.8 |
| **pSer295** | 348.5 ± 60.5 |

#### Supplementary Table 3: C-terminal peptide ligands used for FP assays

| Peptide ligand | Sequence | Calculated MW [M+H$^+$] | Observed MW [M+H$^+$] | Purity UPLC at 214 nm |
|---|---|---|---|---|
| **GluN2A** | TAMRA-NNG-KKMPSIESDV | 1831.8 | 1832.1 | 95.3% |
| **GluN2B** | TAMRA-NNG-YEKLSSIESDV | 1967.9 | 1967.8 | 96.6% |
| **Stargazin** | TAMRA-NNG-NTANRRTTPV | 1827.9 | 1827.9 | 97.7% |
| **Kv1.4** | TAMRA-NNG-SNAKAVETDV | 1731.8 | 1731.9 | 97.6% |
| **Kv1.7** | TAMRA-NNG-PAGKHMVTEV | 1766.8 | 1766.0 | 97.4% |

**Supplementary Table 4: $K_d$ values from FP assays with C-terminal peptide ligands**

All protein variants of PDZ1, PDZ1-2 and PSD-95 were tested for binding to selected C-terminal peptide ligands and the binding constants ($K_d$) in µM calculated. The WT proteins were tested for binding to all five ligands; GluN2A, GluN2B, stargazin, Kv1.4 and Kv1.7. Protein variants phosphorylated on Ser78 (*) and Ser116 (*) are indicated. N.B. = not binding.

| | Protein | GluN2A | GluN2B | Stargazin | Kv1.4 | Kv1.7 |
|---|---|---|---|---|---|---|
| 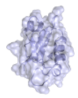 | **PDZ1** | | | | | |
| | WT | 7.9 ± 0.2 | 12.4 ± 0.2 | 27.3 ± 0.8 | 9.7 ± 0.3 | 18.1 ± 0.4 |
| | pSer73 | 13.3 ± 1.1 | 26.3 ± 2.6 | - | - | - |
| | pSer78 | - | N.B. * | N.B. * | - | - |
| | pSer142 | - | - | - | 11.9 ± 0.3 | 21.1 ± 0.6 |
| 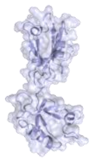 | **PDZ1-2** | | | | | |
| | WT | 1.2 ± 0.01 | 1.9 ± 0.03 | 7.52 ± 0.14 | - | - |
| | pSer73 | 1.55 ± 0.02 | 2.11 ± 0.02 | - | - | - |
| | pSer78 | - | 1.79 ± 0.01 * | 8.97 ± 0.24 * | - | - |
| | pSer116 | - | - | 5.57 ± 0.16 * | - | - |
| 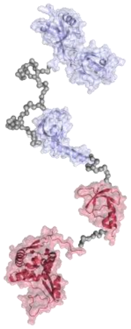 | **PSD-95** | | | | | |
| | WT | 0.83 ± 0.03 | 1.19 ± 0.03 | 2.66 ± 0.07 | 1.27 ± 0.05 | 1.71 ± 0.06 |
| | pSer73 | 1.05 ± 0.02 | 1.23 ± 0.02 | - | - | - |
| | pSer78 | - | 1.34 ± 0.01 * | 3.66 ± 0.02 * | - | - |
| | pSer116 | - | - | 3.2 ± 0.08 * | - | - |
| | pSer142 | - | - | - | 1.41 ± 0.03 | 1.96 ± 0.03 |
| | pSer295 | - | - | 3.55 ± 0.07 | - | - |
